# Resolving isomeric posttranslational modifications using a nanopore

**DOI:** 10.1101/2021.11.28.470241

**Authors:** Tobias Ensslen, Kumar Sarthak, Aleksei Aksimentiev, Jan C. Behrends

**Author notes:** **Correspondence and requests for materials** should be addressed to A.A. or J.C.B.

## Abstract

Posttranslational modifications (PTMs) of proteins are crucial for cellular function but pose analytical problems, especially in distinguishing chemically identical PTMs at different nearby locations within the same protein. Current methods, such as liquid chromatography-tandem mass spectrometry, are technically tantamount to *de novo* protein sequencing^1^. Nanopore techniques may provide a more efficient solution, but applying the concepts of nanopore DNA strand sequencing to proteins still faces fundamental problems^2–4^. Here, we demonstrate the use of an engineered biological nanopore to differentiate positional isomers resulting from acetylation or methylation of histone protein H4, an important PTM target^5,6^. In contrast to strand sequencing, we differentiate positional isomers by recording ionic current modulations resulting from the stochastic entrapment of entire peptides in the pore’s sensing zone, with all residues simultaneously contributing to the electrical signal. Molecular dynamics simulations show that, in this whole-molecule sensing mode, the non-uniform distribution of the electric potential within the nanopore makes the added resistance contributed by a PTM dependent on its precise location on the peptide. Optimization of the pore’s sensitivity in combination with parallel recording and automated and standardized protein fragmentation may thus provide a simple, label-free, high-throughput analytical platform for identification and quantification of PTMs.

---

Bacterial pore-forming proteins reconstituted in synthetic lipid membranes have been used as electrical molecular sensors in a variety of tasks^7,8^. Sequencing of DNA with biological pores, in particular, has become an established technique^9–11^. Generally, in molecular sensing by nanopores, the modulation of a nanopore’s ionic conductance by a single molecule is used to infer that molecule’s identity or sequence. Sequencing by nanopores, as currently done, is based on enzymatically controlled threading of a stretched polymer (DNA or RNA) through pores with short dominant constrictions (MspA, CsgG), so that the partial block of the ionic current is determined by a few monomers (bases) only. By contrast, in whole molecule sensing, all segments of a molecule are simultaneously present within the nanopore and contribute to the current block, which enables the use of a nanopore as a single molecule size spectrometer ^12–19^. Recently, the pore formed by the protein toxin aerolysin (AeL) of *Aeromonas hydrophila* was used in this way for high-resolution discrimination of oligo-PEGs^20^, short oligonucleotides^21^ and oligopeptides^22–24^.

Harnessing of nanopore technology for applications in proteomics has been an active area of research for some time^2,25^, including the detection of PTMs^26–29^. Sequencing proteins by nanopore threading, which would in principle allow detection of PTMs in a site-specific manner, is presently in early stages of development, but still faces the lack of processive motors suitable for moving a polypeptide chain through the nanopore and of physical mechanisms for keeping an unevenly charged peptide taut through the nanopore’s constriction^3,4,25,30^. Here we show that site-specific detection of histone PTMs is feasible by whole-molecule nanopore sensing of peptide fragments, requiring neither sequential threading nor uniform charge of the analyte. In contrast to recent work using engineered electrostatic interactions to discriminate phosphoserine and phosphothreonine residues^27^, our strategy is not limited to charge-conferring modifications and is capable of differentiating true positional isomers. Furthermore, our all-atom molecular dynamics (MD) simulations^31,32^ show that the ability to determine the sites of chemically identical modifications originates from an analyte-induced modulation of the nonuniform electric field inside the nanopore, suggesting a universal approach for electrical sensing of molecular shape.

For our proof-of-principle experiments, we chose a decapeptide sequence occurring near the N-terminus of histone protein 4, H4 (K8-R17) (**Fig. 1a**). This protein fragment, H4f., contains three lysine residues (K8, K12, K16) at which acetylation has important functional consequences for gene expression^32,33^. We chose the sequence to include, in particular, lysine K16, as its acetylation was recently found to be important for human development^34^, as well as to be an intergenerational gene activation signal in Drosophila^35^. We chose the sequence to be shorter than the common N-terminal tryptic fragment H4(G4-R17) to be commensurate with the length of peptides known to provide good size resolution with the aerolysin pore^22,23^.

**Fig. 1.**
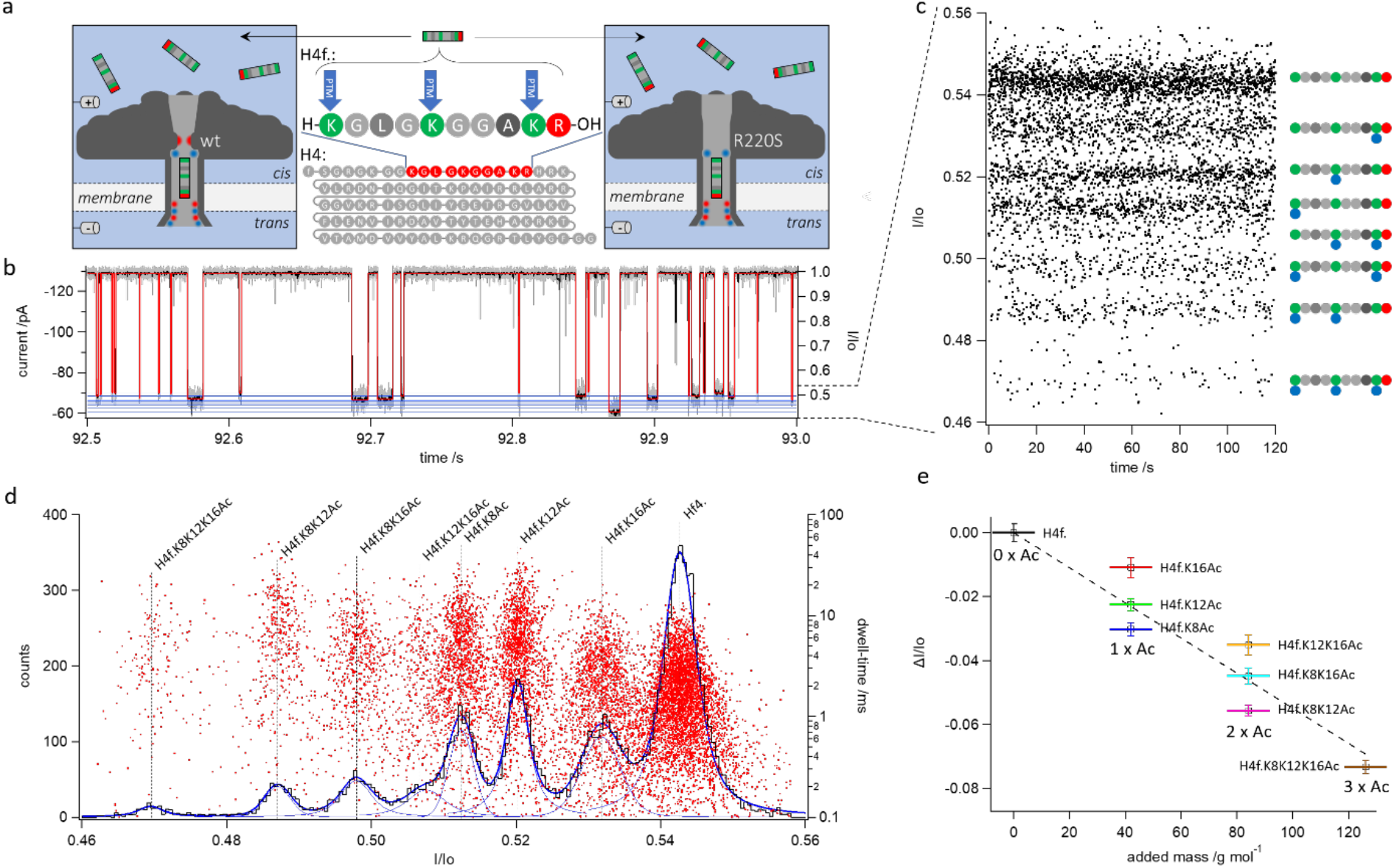
Discrimination of acetylation-derived positional isomers with the R220S mutant of the aerolysin pore. **a**, Experimental set-up and peptide constructs used. Left: schematic representation of the wt-AeL pore reconstituted in a lipid bilayer membrane with the bulky, positively charged arginine in position 220 (R220) as part of the cis-side constriction. Center: Schematic of the full length human H4 protein, and the fragment (H4f.) used in this study. Blue arrows indicate the positions of the three lysins (K8, K12, K16) susceptible to posttranslational modification by acetylation or methylation. Right: the mutant aerolysin pore in which R220 was replaced by a serine, neutralizing the charge and reducing the extent of the constriction. In both representations of the pore, a peptide is schematically shown residing in the main beta-barrel, the presumptive sensing region. **b**, Half-a-second current trace of an experiment recorded using the R220S pore in the simultaneous presence of H4f, H4f.K8Ac, H4f.K12Ac, H4f.K16Ac, H4f.K8K12Ac, H4f.K8K16Ac, H4f.K12K16Ac and H4f.K8K12K16Ac and digitally postfiltered at 25 kHz (grey) or 2.5 kHz (black) 3 dB cut-off frequency (see Methods). Red trace indicates the output of the detection algorithm (see methods); blue lines indicate the means of 8 preferred current amplitudes in the blocked state corresponding to the 8 peptoforms. Left y-axis indicates the absolute current (pA), right y-axis shows blocked state current normalized by open pore current (I/Io); **c**: scatterplot of I/Io values against time over 2 min of recording. Note the concentration of data points around 8 distinct levels which can be assigned to the various peptoforms (right) by sequential addition (Extended Data Fig. 2). **d**, Black cityscape: histogram of I/Io levels from the experiment shown in b, c, fitted with the sum (thick blue line) of 8 Voigt peaks ^31^ (thin blue lines) corresponding to each probability maximum. Red dots: superimposed scatterplot of resistive pulse dwell-time vs. I/Io value. **e**, Shift of maxima in d with respect to the unmodified H4f.-peptide (black) plotted against mass added by acetylations. Error bars give full width at half maximum of Voigt peaks. Note that despite a linear overall dependence of I/Io on mass, the maxima corresponding to peptoforms of identical mass are clearly separated.

A major consideration for whole-molecule sensing is the dwell-time of the analyte in the nanopore. In the presence of noise, longer current blocks give more precise estimation of the relative residual current and thereby, enhance resolution of subtle differences between analyte molecules^8,20,23^ and site-directed mutagenesis of presumptive pore-lining residues can be used to prolong analyte interaction with biological nanopores^28,36–38^. On the other hand, measurement error decreases only with the square root of dwell-times and saturation of the pore may become an issue. In exploratory experiments using arginine-based peptides, we identified a novel functional aerolysin mutant, R220S, in which the dwell-times are approximately doubled with respect to the wild-type and which enhanced differentiation of peptides containing single amino acid substitutions (**Extended Data Fig. 1**). We, therefore, used the R220S mutant for the task of differentiating positional peptide isomers.

**Fig. 1** reports results from a typical recording using the R220S aerolysin mutant in the presence of all seven peptide variants (peptoforms), resulting from acetylation of either one, two or all three lysine residues K8, K12 and/or K16 (H4f.K8Ac, H4f.K12Ac, H4f.K16Ac, H4f.K8K12Ac, H4f.K8K16Ac, H4f.K12K16Ac and H4f.K8K12K16Ac), as well as the unmodified fragment Hf4. As shown in **Fig. 1**b, the open pore current of approximately −130 pA at a driving force of −60 mV is, at irregular intervals, interrupted by blocks that reach various, reduced current levels which are situated around 50 % of the open pore current Io (right axis: relative residual current I/Io≅0.5). A plot of the computed event-averaged mean current values of these blocks vs. time (**Fig. 1**c) reveals them to be concentrated at eight preferred relative residual current values. A histogram showing the corresponding probability maxima is shown in **Fig. 1**d. These levels were unequivocally assigned to each species by sequential addition of the peptoforms (see **Extended Data Fig. 2**). Surprisingly, as also shown by the multi-Voigt peak fits^31^ in **Fig. 1**d and **Extended Data Fig. 2**, there is good to excellent separation between the maxima corresponding to the positional isomers for single acetylation (H4f.K16Ac, H4f.K12Ac, H4f.K8Ac) and double acetylation (H4f.K12K16Ac, H4f.K8K16Ac, H4f.K8K12Ac). As expected, the sequence of I/Io-values follows mass, or more likely global volume^8,23^, so that increased overall acetylation induces a shift towards deeper blocks. Indeed, a plot of the shift of the mean relative residual current value (ΔI/Io) with respect to unmodified H4f. vs. the mass added by acetylation shows an overall linear relationship (**Fig. 1**e). However, the peptoforms of equal mass (positional isomers) are well separated from each other (**Fig. 1**e), with acetylations towards the N-terminus providing deeper blocks (**Extended Data Fig. 3**a, b). Note the reduction of peak area with increased acetylation due to neutralization of the positive charges of the lysines and the resulting weakening of electrophoretic force on the analyte (Extended Data Fig. 3c).

The clear separation between the population maxima is mainly possible because all peptoforms induce blocks with characteristic dwell-times well above 1 ms in the R220S variant (see scatterplot in **Fig. 1**d and **Extended Data Fig. 3**d and **Extended Data Fig. 4**), with acetylation propitiously increasing these values, thus allowing efficient reduction in measurement error by digital averaging for all species. This contrasts with the wild type aerolysin pore, where acetylation induces a strong reduction in dwell-times to well below 1 ms (Extended Data Fig. 5 - **Extended Data Fig. 8**). As evidenced by sequential addition experiments, the different peptoforms block the ionic current to unique levels also in the wild-type pore, and also in the same sequence. However, the measurement error due to shorter dwell-times produces considerable overlap between the I/Io-probability maxima for peptoforms of equal mass (Extended Data Fig. 6, Extended Data Fig. 8a, b).

In essence, our findings show that, in both wild type and mutant pores, chemical modifications of residues affect the residual current in a manner dependent on the position of the modification in the peptide chain. This result opens a range of new possibilities for the use of nanopores in probing the structure of single molecules and would not be predicted in the framework of the constant field assumption for ion transport^39^.

To determine the molecular mechanism enabling the site-selective differentiation of PTMs, we used the steered MD (SMD) protocol^40,41^ to pull an H4f.K8Ac peptide through the transmembrane pore of the aerolysin variant R220S (**Fig. 2**a). The sequence of microscopic conformations generated by the SMD run was analyzed using the steric exclusion model (SEM)^42^ to determine the relative residual current (I/Io) as a function of the peptide’s location within the pore, **Fig. 2**b. The residual current has its highest value close to 1.0 as the peptide enters the pore from the *cis*-side and the lowest value of 0.1 as it passes through the *trans*-side constriction of aerolysin. Starting from the ensemble of conformations generated by pulling the H4f.K8Ac peptide, we obtained conformational ensembles for H4f., singly (H4f.K8Ac, H4f.K12Ac, H4f.K16Ac), doubly (H4f.K8K12Ac, H4f.K8K16Ac, H4f.K12K16Ac) and triply (H4f.K8K12K16Ac) acetylated peptides by computationally mutating the side chains of the respective acetylated residues^23^. Repeating the SEM calculations for the modified peptides produced seven additional profiles of the relative residual current, **Fig. 2**b, all of which appear to be nearly indistinguishable from that of H4f.K8Ac, except in certain regions of the pore. The average simulated residual current of all eight peptides, **Fig. 2**c, was found to be in good agreement with the experimental values when the currents were averaged over the complete sensing volume of wt-AeL^23^, or over the region where the deviations were observed (**Extended Data Fig. 9**). From repeated SMD simulations we learned, however, that the sequence of blockade level’s ordering does depend on the global conformation of the peptides, in particular on the global orientation of the peptide with respect to the nanopore (see **Extended Data Fig. 10** and **Extended Data Fig. 11**).

**Fig. 2.**
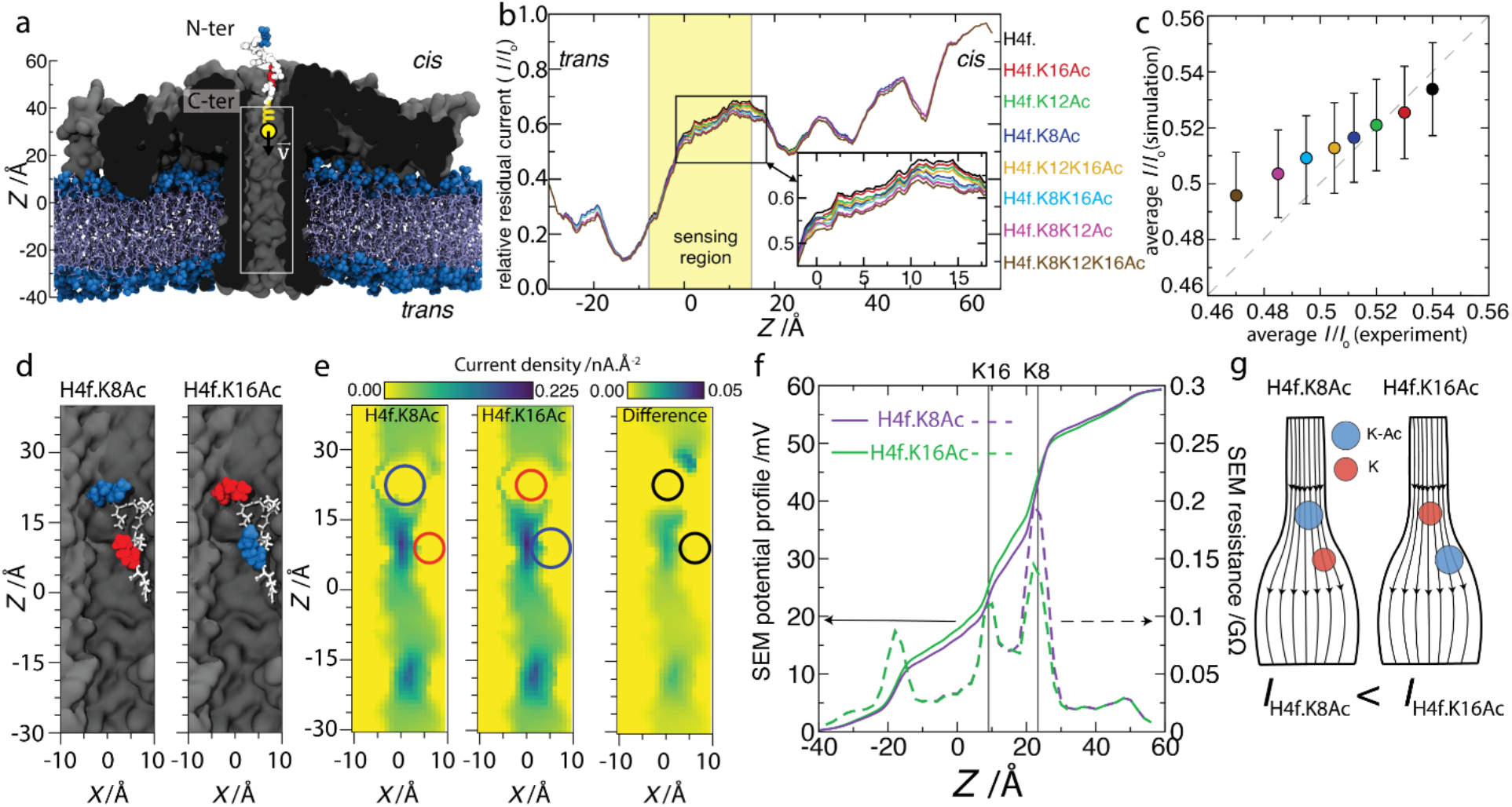
Microscopic mechanism of site-selective PTM detection. **a**, Initial state of a 100 ns steered MD simulation where an H4f.K8Ac peptide (vdw spheres) is pulled by a harmonic spring with a constant velocity of 1 Å/ns through an aerolysin nanopore (cutaway molecular surface), embedded in a lipid membrane (blue) and submerged in 2 M KCl electrolyte (not shown). The C-terminus of the peptide is oriented towards the trans-side of the membrane. The white rectangle indicates the location of the nanopore volume shown in panels d and e. **b**, Relative residual current versus center of mass z coordinate of the acetylated H4f. peptides. The coordinate axis is defined in panel a. The currents were computed using SEM^42^ and running-averaged with a 5 Å window. The highlighted region shows the sensing volume (−9 Å < z < 15 Å) of the wt-AeL pore^23^. The inset shows a zoomed-in view of the main plot. **c**, Simulated vs. experimental average relative currents produced by the acetylated peptides within the sensing region of aerolysin. Colors represent peptide variants as defined in panel b. The error bars show the standard error calculated using 100 current values from the sensing region of the nanopore. **d**, Representative conformation of the H4f. K8Ac peptide in the sensing region of the nanopore and the corresponding computational model of the H4f.K16Ac peptide. The peptide residues are drawn as white sticks except residues K8 and K16, which are shown as vdW spheres colored according to their acetylation state: acetylated (blue) and unmodified (red). **e**, local density of transmembrane ion current (its z component) near the H4f. K8Ac (left) and H4f. K16Ac (center) peptides and their difference (H4f.K16Ac-H4f.K8Ac, right). The currents were computed using SEM over a 1 Å grid. The heat map shows a cross section of the nanopore volume along the pore axis. The circles indicate the approximate locations of the K8 and K16 residues. **f**, Electrostatic potential along the nanopore axis for the H4f.K8Ac (purple) and H4f.K16Ac (green) peptide systems (solid lines, left axis) and local resistance of the nanopore volume (dashed lines, right axis). The local resistance was computed from the local electrostatic potential using 4 Å segments. **g**, Schematics illustrating the microscopic mechanism enabling site-specific PTM detection via a blockade current measurement. The blue and red balls represent acetylated and unmodified lysine residues, respectively.

Having established a computational model that reproduces experimentally measured differences in the blockade current, we identified the molecular mechanism that enables site-specific determination of PTMs. Whereas a current difference produced by either acetylation or deacetylation is expected, as such a modification changes the overall volume excluded by the peptide from the conductive volume of the nanopore, the transfer of the same modification from one site to another does not alter the excluded volume and thus any effect on pore conductance requires a more detailed examination. To this end, we compared the local density of transmembrane ionic current (the current’s z component) in the nanopore blocked by either H4f.K8Ac or H4f.K16Ac peptides, which molecular configurations differ only by the site of acetylation, **Fig. 2**d (see Extended Data Fig. 12 for similar analysis of another peptide conformation from the same MD trajectory). The local current density map, **Fig. 2**e, shows low currents in the regions occupied by the N-terminus of the peptide (i.e., around z=22 Å, residue 8 and 9) and high currents in the parts of the nanopore devoid of the peptide atoms (i.e., around z=6 Å, residues 15-17). The electrostatic potential profile computed for the two peptide conformations, **Fig. 2**f, reveals that the sharpest drop of the potential occurs in the vicinity of residue K8, and that acetylation of that residue considerably increases resistance of the adjacent nanopore volume (**Fig. 2**f, dashed lines). In contrast, acetylation of residue K16 introduces negligible changes to the local nanopore resistance. This is in line with the experimental finding which shows a deepening of the block of the pore as the lysines towards the N-terminus of the peptide are acetylated (Extended Data Fig. 3b). Thus, our analysis points to a microscopic mechanism where a peptide’s confinement within the nanopore creates a conductive volume of a highly non-uniform shape. Because of this non-uniform shape, a local change of the volume’s cross section due to a PTM can alter the overall conductance of the nanopore by an amount that depends on the location of the PTM within the conductive volume, similar to an effect produced by placing a boulder in either rapids or a backwater section of a mountain stream, **Fig. 2**g.

To determine whether our result can, in principle, be generalized to other PTMs of histone proteins, we attempted to analyze site-selective mono- and trimethylation of lysines K8, K12 and K16 using the R220S mutant aerolysin pore. In order to allow direct comparison with the acetylation results, we used the same H4f. as above, despite the fact that K20, rather than K8, K12 and K16, is reported to be a known H4 methylation target^43,44^.

As shown in **Fig. 3**a, clearly different maxima were apparent in the relative residual current (I/Io) histogram for nonmethylated and singly, doubly and triply monomethylated forms of H4f. As in the case of acetylation, successive monomethylation at one, two or three residues resulted in a progressive shift of the I/Io-maxima towards deeper blocked levels that was linearly related to added mass (**Fig. 3**d). In keeping within the smaller added mass of methylation (14 g/mol) vs. acetylation (42 g/mol), the total shift by triple monomethylation was less than that by triple acetylation; however, the steepness of the relationship between the shift of ΔI/Io and the added mass was higher for methylation than for acetylation. While the histogram in **Fig. 3**a shows no obvious separation between the maxima for the positional isomers H4f.K8me, H4f.K12me, H4f.K16me or H4f.K8K12me, H4f.K8K16me, H4f.K12K16me, it became evident in sequential addition experiments that each species contributed a population of resistive pulses with a characteristic mean relative current I/Io.

**Fig. 3.**
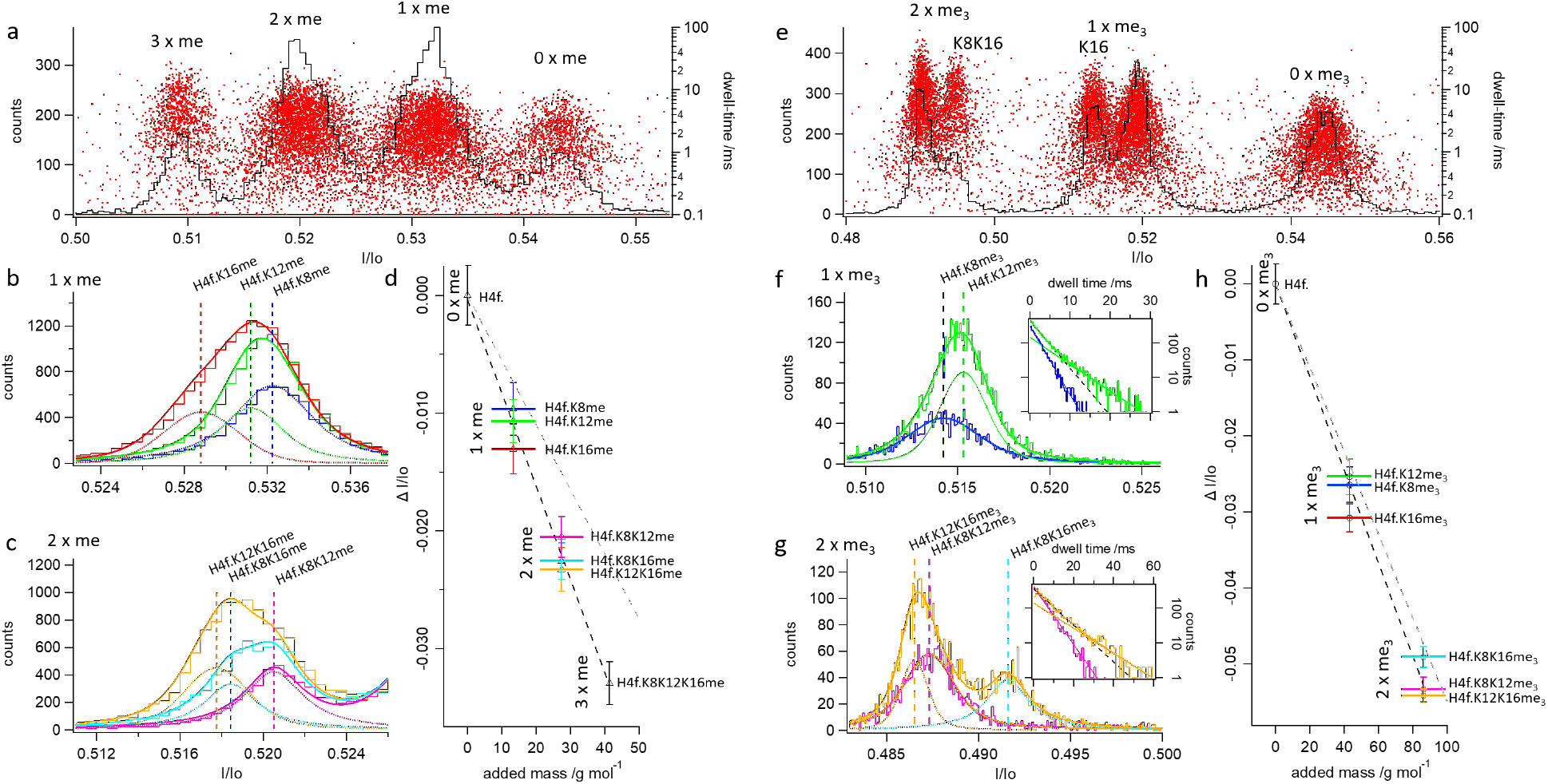
Discrimination of positional isomers from mono- and trimethylation with the R220S mutant. **a**, Histogram of I/Io-values (black) and scatterplot of dwell-time vs. I/Io (red) in the presence of all singly, doubly and triply monomethylated species, as well as of unmodified H4f. Unlike for acetylation, the positional isomers (singly monomethylated H4f.K8me, H4f.K12me and H4f.K16me, as well as doubly monomethylated H4f.K8K12me, H4f.K8K16me and H4f.K12K16me) are not obviously separated in terms of blockade levels, as their maxima overlap (for assignment of peaks by sequential addition see **Extended Data Fig. 13** - **Extended Data Fig. 15**). However, as shown in b and c, sequential addition of isomeric species leads to successive broadening of the maxima, which can be accounted for by the contribution of populations isomeric species with different means. By fitting the I/Io-histograms (cityscapes) with a sum (thick lines) of Voigt peaks (thin lines), the respective position of the maximum of the added peptide can be determined. **b**, Sequential addition of singly monomethylated species H4f.K8me (blue), after addition of H4f.K12me (green) and after addition of H4f.K16me (red); **c**: sequential addition of doubly monomethylated species H4f.K8K12me (magenta), after addition of H4f.K8K16me (turquois) and after addition of H4f.K12K16me (gold); For a complete account of the sequential addition experiment see **Extended Data Fig. 15**. **d**, Shift of maxima with respect to that for unmodified H4f. plotted against the mass added by monomethylation; **e**: histogram of I/Io-values (black) and scatterplot of dwell-time vs. I/Io (red) in the presence of unmodified H4f., as well as all singly and doubly trimethylated species. In this case, the maxima for H4f.K8me_3_ and H4f.K12me_3_, as well as those for H4f.K8K12me_3_ and H4f.K12K16me_3_, respectively, overlap in one peak whereas those for H4f.K16me_3_ and H4f.K8K16me_3_ are strongly shifted to deeper (H4f.K16me_3_) and more shallow blocks (H4f.K8K16me_3_). For assignment of peaks by sequential addition see **Extended Data Fig. 16** - **Extended Data Fig. 18**; **f**: sequential addition of singly trimethylated species H4f. K8me_3_ (blue) and after addition of H4f.K12me_3_ (green). Note the shift towards longer dwell-times and failure of monoexponential fit (dashed line) after addition of H4f.K12me_3_; **g**, Sequential addition of singly trimethylated species H4f.K8K12me_3_ (turquois), after addition of H4f.K8K16me_3_ (magenta) and H4f.K12K16me_3_ (gold). Insets in f and g: dwell-time histograms for the two overlapping species. Note the shift to longer dwell-times and departure from monoexponential distribution (dashed lines) upon addition of H4f.K12me3 or H4f.K12K16me_3_. **h**, Shift of maxima with respect to that for unmodified H4f. plotted against the mass added by trimethylation; All histograms in b, c and f, g were generated from recordings lasting 300 s, making them directly comparable. Grey dashed lines in e and h represent the relationship between the shift in I/Io and added mass for acetylation (see **Fig. 1**e).

As shown in **Fig. 3**b, equimolar addition of H4f.K12me to the *cis*-compartment already containing H4f.K8me not only made the blockade events more frequent, but also broadened the I/Io-distribution and shifted its maximum towards a smaller value. A similar trend was observed when H4f.K16me was added to a *cis*-compartment containing H4f.K8me and H4f.K12me. In both cases, this shift and broadening could be accounted for by the contribution of an additional population, characterized by a slightly shifted mean, as shown by the Voigt fits in **Fig. 3**b and **Extended Data Fig. 15**a-c. Different population maxima could also be determined by sequential addition of H4f.K8K12me, H4f.K8K16me and H4f.K12K16me (**Fig. 3**c and **Extended Data Fig. 15**d-f).

We note that, while not allowing for single-passage identification, as for acetylation, the site-specific sensitivity of the mutant pore for monomethylation is sufficient to allow for a quantitative analysis of methylated peptoforms, as shown above, given appropriate calibration of on-rates to correct for the loss of electrophoretic susceptibility.

The study of lysine trimethylation of H4f. suggests that relative abundance of isomer populations can not only be discerned from the blockade levels (I/Io) but also from the analysis of the dwell-time distributions. The positional isomers H4f.K8me_3_ and H4f.K12me_3_ could not be *prima facie* distinguished on the basis of the I/Io-distributions in the presence of all seven species but H4f.K16me_3_ showed a clearly distinct maximum that was shifted towards lower residual currents. Conversely, the maxima for doubly trimethylated H4f.K8K12me_3_ and H4f.K12K16me_3_ were lumped together in one peak, while H4f.K8K16me_3_ produced a distinct peak, shifted towards higher residual current levels (**Fig. 3**, **Extended Data Fig. 16**). As before, closer analysis of sequential addition experiments revealed a small shift in the population mean upon adding H4f.K12me_3_ to H4f.K8me_3_ or H4f.K12K16me_3_ to H4f.K8K12me_3_, which enabled the use of the fitting procedure previously shown for monomethylation to extract separate I/Io values for the two overlapping species (**Fig. 3**f, g and **Extended Data Fig. 18**). Furthermore, by addition of H4f.K12me_3_ to H4f.K8me_3_ or H4f.K12K16me_3_ to H4f.K8K12me_3_ the monoexponential distribution of event dwell-times contained in the maximum (τ≈2 and 5 ms, respectively) was converted into a double-exponential distribution due to the appearance of a second exponential population with longer dwell-times (τ’≈5 and 10 ms, respectively, **Fig. 3**f, g inset and **Extended Data Fig. 17**). The relative weights of the two exponential components may, therefore, also be used to estimate the relative abundance of these isomeric peptoforms following appropriate correction for on-rates. Again, we found an overall linear relationship between the shift in I/Io, added mass and the steepness for the relation for trimethylation was indistinguishable from that for acetylation. Synthesis of triply trimethylated H4f. was attempted at two commercial sources, but failed.

In summary, we have shown that an engineered aerolysin nanopore can be used to quantitatively analyze both acetylation and methylation in a position-sensitive fashion by distinguishing between peptoforms of equal mass on the basis of depth-of-block and, in some cases, duration of resistive pulses. The mechanism of discrimination by resistive current can be explained by the action of a strongly inhomogeneous electric field that is shaped by the pore structure, as well as by the conformation of the confined peptide itself (**Fig. 2**). The dependence on the peptide’s conformation also explains for the fact that the sequence of mean I/Io-values for PTM of the different lysines is different for each of the three PTMs analyzed, see **Extended Data Fig. 19**. The higher sensitivity of the mutant pore relative to the wild-type likely originates from a powerful molecular trap that the aerolysin pore provides^23^, and which becomes more efficient upon removal of the bulky and positively charged R220 (**Extended Data Fig. 20**, **Extended Data Fig. 21**). Additional mutations of the pore can be envisaged which, in combination with R220S, will lead to further kinetic optimization also for the use with tryptic peptides containing more than ten amino acids. To further increase differentiation among the positional isoforms, future work will focus on reducing instrumental noise^45^ and on protein engineering to reducing the conformational degree of freedom of peptides in the pore. Thus, in a combination of parallel high-resolution electrophysiology^15,18^ with standardized and automated enzymatic generation of peptide fragments^46^, whole molecule sensing with the aerolysin nanopore has the potential of being implemented in a practical technology to rapidly detect PTMs with positional resolution without requiring strand sequencing of proteins.

## Methods (online only)

### Mutagenesis, Expression and Purification

AeL-R220S single point mutagenesis was carried out following a four primer PCR protocol, followed by standard overlap-PCR. Gene flanking and mutagenesis primers were designed with the sequences depicted below and purchased from Eurofins Genomics GmbH (Ebersberg, Germany).

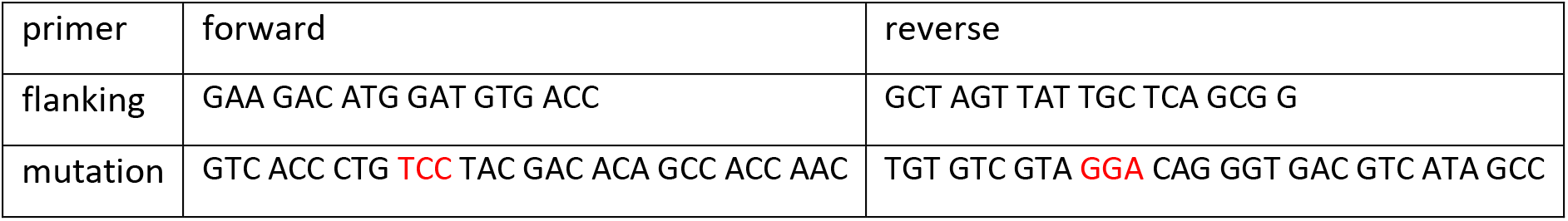

The mutated insert, as well as the pET22b(+) vector were restricted by BamHI and XhoI (Thermo Fisher Scientific, Waltham, MA, USA) and ligated with T4 DNA Ligase (Fermentas - Thermo Fisher Scientific, Waltham, MA, USA). Mini-scale plasmid preparation was performed after transformation of XL1-Blue competent cells via heat-shock prior to DNA sequencing (Eurofins Genomics GmbH, Ebersberg, Germany). Subsequent max-scale plasmid preparation was performed after transformation of XL1-Blue competent cells with the pET22b(+)::plb::pAeL-R220S-His6 vector via heat-shock. Successful transformation of the cells with vector material containing the insert was verified by enzymatic control restriction using NdeI, XhoI and BamHI (Thermo Fisher Scientific, Waltham, MA, USA) prior to verification of the final plasmid’s nucleotide sequence (Eurofins Genomics GmbH, Ebersberg, Germany).

Biosynthesis of wt-pAeL and pAeL-R220S was carried out in BL21(DE3)pLysS competent cells after transformation with the corresponding pET22b(+)::plb::pAeL-His6 construct and cultivation at 37 °C. Gene expression was induced at OD600=0.6 by addition of 500 μM IPTG and verified by SDS-PAGE analysis. Cells were further incubated at 18 °C and harvested by centrifugation (1 h, 4400 rpm, 4 °C) when no further change of the OD600 was observed. Cells were washed with cold PBS and pelleted by centrifugation (45 min, 4400 rpm, 4 °C). Cells were disrupted by ultrasonication and debris removed by ultracentrifugation (100’000 xg, 2 h, 4 °C). Protein purification was carried out by Ni-affinity-chromatography (AFI), followed by size-exclusion-chromatography (SE) using an ÄKTA HPLC purifier system (GE Healthcare/Amersham Bioscience, Chalfont St Giles, Great Britain), equipped with a 5 mL HisTrap^™^HP or a Superdex 200-Increase 10/300 GL column. Protein containing AFI elutions were pooled and concentrated to a final volume of 250 μL, prior to SE using Amicon^®^Ultra 15 mL centrifuge filters with a molecular weight cutoff of 10 kDa (Merck KGaA, Darmstadt, Germany). Purity of the SE elutions was verified by SDS-PAGE and the protein concentration determined in a Bradford assay (Coomassie brilliant blue G-250, BioRad, Hercules, CA, USA). Final protein samples were adjusted to 1 mg/mL, nitrogen shock-frozen and stored at −80 °C. Activation of pAeL was carried out before an experiment by application of trypsin (Promega, Madison, WI, USA) at room-temperature within 15 min.

### Electrophysiology and data analysis

Nanopore recordings were performed using a modified version of the Orbit16 device (Nanion Technologies, Munich, Germany using MECA 16 microelectrode cavity arrays (Ionera Technologies, Freiburg, Germany) as previously detailed^8,18,23^. Briefly, one of 16 coplanar gold lines leading to MECA 16-cavities (50 μm diam.) was directly contacted with a 1 cm unshielded silver wire to the head-stage of an Axopatch 200B (Molecular Devices, Sunnyvale, CA, USA) patch clamp amplifier operated in capacitive feedback mode at 50 mV/pA gain with the internal low-pass-filter set to 100 kHz cut-off (−3dB). The output signal was passed through an external low-pass Bessel filter (npi electronic, Tamm, Germany, 8 pole, custom version of LHBF-48X-8HL) set to 50 kHz cut-off and digitized at 1 MHz sampling rate using a PCI-6251 16 bit ADC interface (National Instruments, Austin, TX, USA) controlled by GePulse software (Michael Pusch, University of Genoa, Italy). Event-averaged current level detection was performed using thresholding on the differentially rectified signal as shown before^20,23^ and explained in detail in Ref^8^. Further analysis and graphing were performed using IgorPro8 (Wavemetrics, Portland, OR, USA).

### Molecular dynamics simulations

#### General MD methods

all MD simulations were carried out using NAMD2^47^, a 2 fs integration timestep, periodic boundary conditions, the CHARMM36^48^ force field and CUFIX corrections to non-bonded interactions^49^. The covalent bonds containing hydrogen atoms in water were restrained using the SETTLE algorithm^50^, and in protein using the RATTLE algorithm^51^ Long-range electrostatic interactions were evaluated using the particle mesh Ewald method^52^ on a 1 Å grid. Constant pressure was maintained using a Nosé-Hoover Langevin piston^53^. Constant temperature was maintained by coupling the non-hydrogen atoms of the lipid to a Langevin thermostat (Brünger, A. T. X-PLOR: version 3.1: a system for x-ray crystallography and NMR (Yale University Press, 1992)) with a 5 ps^−1^ damping constant. The van der Waals forces were calculated with a switching distance of 10 Å and a cutoff of 12 Å. Local interactions were evaluated every time step and full electrostatics every second time step.

#### All-atom model of aerolysin

the all-atom model of the aerolysin-R220S nanopore was built starting from a pre-equilibrated (20 ns), solvated model of wt-AeL embedded in a 1,2-diphytanoyl-sn-glycero-3-phosphocholine (DPhPC) ¬lipid bilayer^23^ The R220S mutation was introduced using the psfgen package of VMD^54^ The number of K^+^ and Cl^−^ ions was adjusted to produce an electrically neutral system containing 2 M KCl solution. Upon assembly, the system underwent 1’000 steps of energy minimization and then equilibrated in the constant number of atoms (430’567), pressure (1 bar) and temperature (293 K) ensemble for 5 ns while restraining the C-α atoms of the protein to their initial coordinates. During the equilibration, the area of the lipid bilayer was kept constant while the system changed its size along the bilayer normal. All subsequent simulations were carried out in a constant number of atoms, volume, and temperature ensemble, where the volume (18.93 nm × 18.93 nm × 11.56 nm) was set to the average value observed within the 5 ns equilibration. The transmembrane bias V=-60 mV was induced by applying a constant electric field, E=−V/Lz, normal to the lipid bilayer, where Lz was the dimension of the system along normal to the bilayer^55^ Each C-α atom of aerolysin was harmonically restrained (with a spring constant of 69.5 pN/nm) to maintain the same coordinate as in the last frame of the equilibration trajectory.

#### Simulations of current blockades

an all-atom model of the H4f. peptide (H-KGLGKGGAKR-OH) was obtained by mutating the side chains of a pre-equilibrated 10-residue peptide taken from a previous study^56^. The N-terminal lysine was acetylated to obtain a microscopic model of the H4f.K8Ac peptide. The peptide was then aligned with the nanopore axis and placed near the *cis*-entrance of the pore such that the C-terminal of the peptide faced the *trans*-side of the pore (**Fig. 2**a). The peptide was then pulled along the pore axis using the SMD method^40,41^, while restraining the C-α atoms of aerolysin to their equilibrated coordinates. During the 100 ns SMD run, the center of mass (CoM) of the peptide was coupled to a template particle by means of a harmonic potential (with a spring constant of 4’865 pN/nm) while the templated particle was pulled along the nanopore axis with a constant velocity of 1 Å/ns. The residual current was then calculated using SEM^42^ for a set of microscopic configurations taken from the SMD simulation every 0.2 ns, yielding 500 relative residual current values. The currents were sorted in ascending order according to the CoM z coordinate and then averaged using a 5 Å running average window. These averaged currents are plotted in **Fig. 2**b. Using our previously described method ^23^, each conformation was computationally mutated to obtain microscopic representations of H4f. and all single (H4f.K8Ac, H4f.K12Ac, H4f.K16Ac), double (H4f.K8K12Ac, H4f.K8K16Ac, H4f.K12K16Ac) and triple (H4f.K8K12K16Ac) acetylation peptides. The residual currents for each mutated peptide conformations, **Fig. 2**b, were calculated and averaged using the same protocol.

## Acknowledgements

TE was partly funded by a PhD fellowship in the framework of the International Graduate College 1642 “Soft Matter Science: Concepts for the Design of Functional Materials” of the Deutsche Forschungsgemeinschaft (DFG). Work in JCB’s laboratory was funded by the German Ministry for Research and Education through Project Management PTJ (TseNareo), by the BW Foundation through Project Management VDI (MSDS-BioMem) and by the Ministry of Commerce of the State of Baden-Württemberg in the Framework of the Forum Gesundheitsstandort Baden-Württemberg (TechPatNano). K.S. and A.A. acknowledge support from the US National Science Foundation (PHY-1430124). The supercomputer time was provided through the XSEDE Allocation Grant MCA05S028 and the Leadership Resource Allocation MCB20012 on Frontera of the Texas Advanced Computing Center. We thank Dr. Gerhard Mittler, Max Planck Institute of Immunobiology and Epigenetics, Freiburg, for helpful suggestions.

## Author contributions

T.E. and J.C.B. conceived and designed experimental work, T.E. performed electrophysiological experiments, molecular biology and protein purification, T.E. and J.C.B. analysed experimental data, K.S. and A.A. designed MD simulations, K.S. performed MD simulations, K.S. and A.A. analysed MD simulations, T.E. J.C.B. K.S. and A.A prepared figures, T.E., J.C.B. and A.A. wrote the manuscript.

## Extended Data Figures

**Extended Data Fig. 1.**
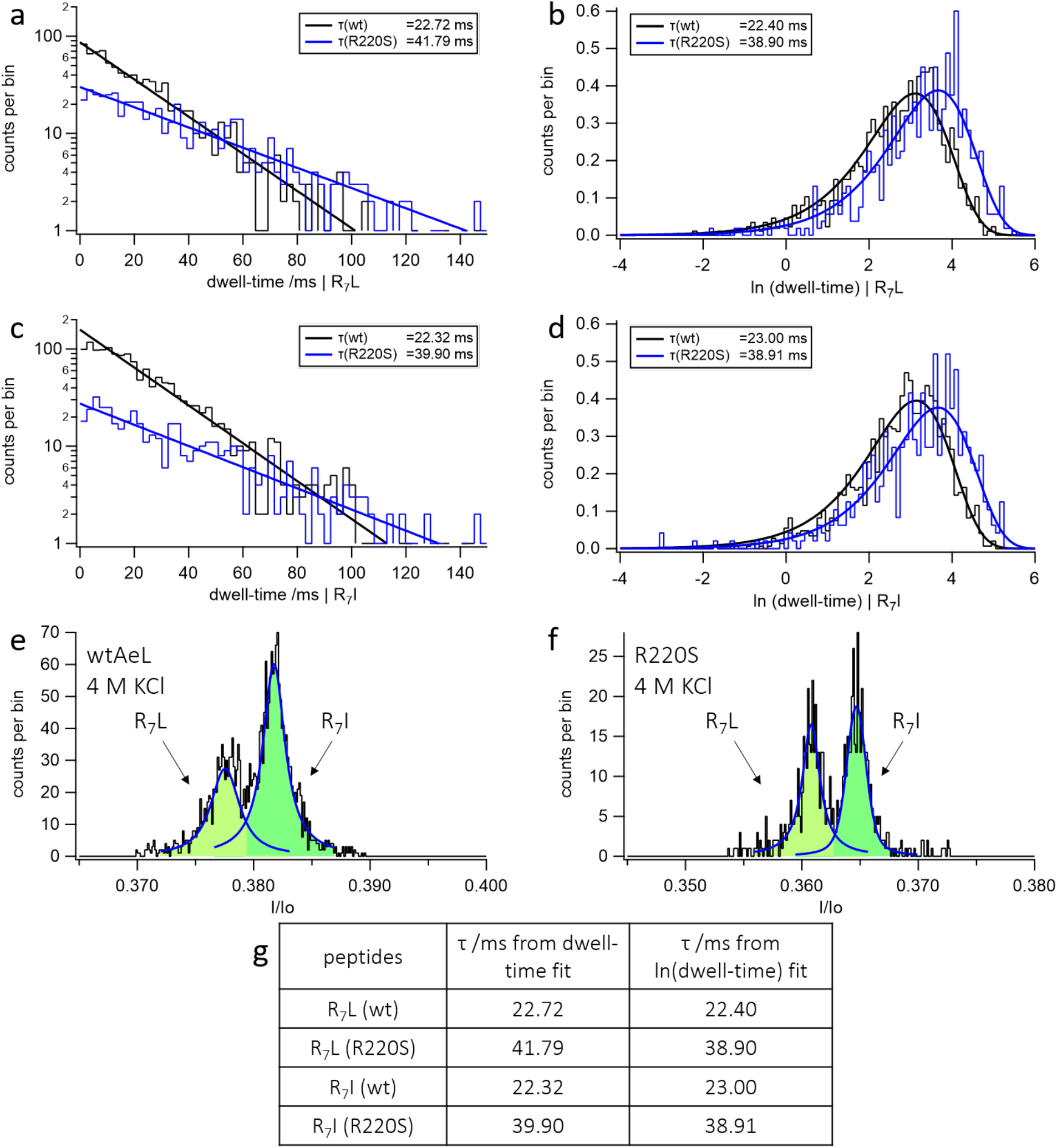
Dwell-time prologation due to R220 single point mutagenesis. **a-d**, Prolongation of dwell-times of hepta-arginine-leucin, R_7_L, and hepta-arginine-isoleucin, R_7_I, peptides in the R220S mutant (blue) vs. wt-AeL pore (black). **e, f**, enhanced discrimination of the isomeric peptides using the R220S mutant.

**Extended Data Fig. 2.**
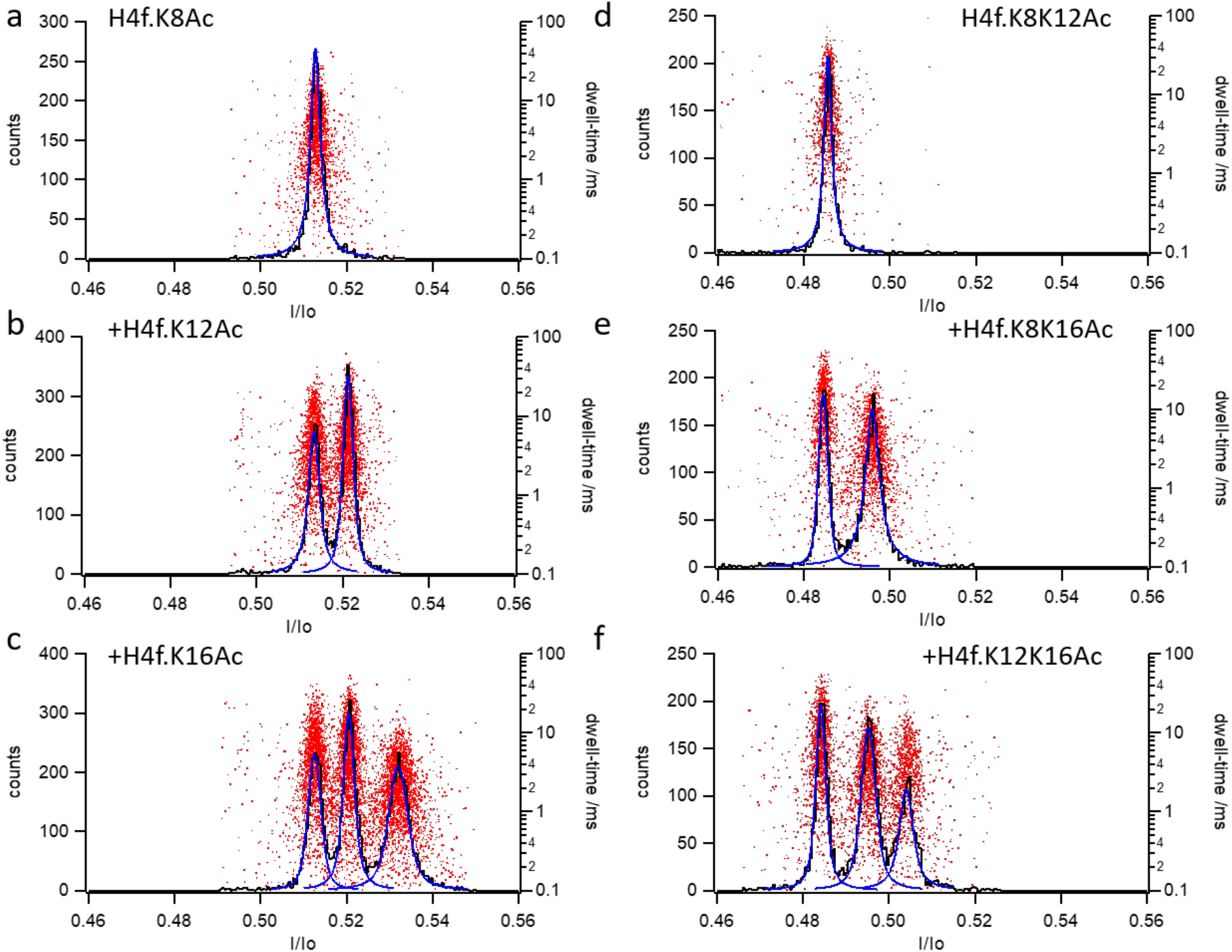
Assignment of maxima for acetylated species of Hf4. by sequential addition of H4f. peptides with one (**a-c**) or two (**d-f**) acetylated residues using sequential addition with the R220S mutant pore.

**Extended Data Fig.3.**
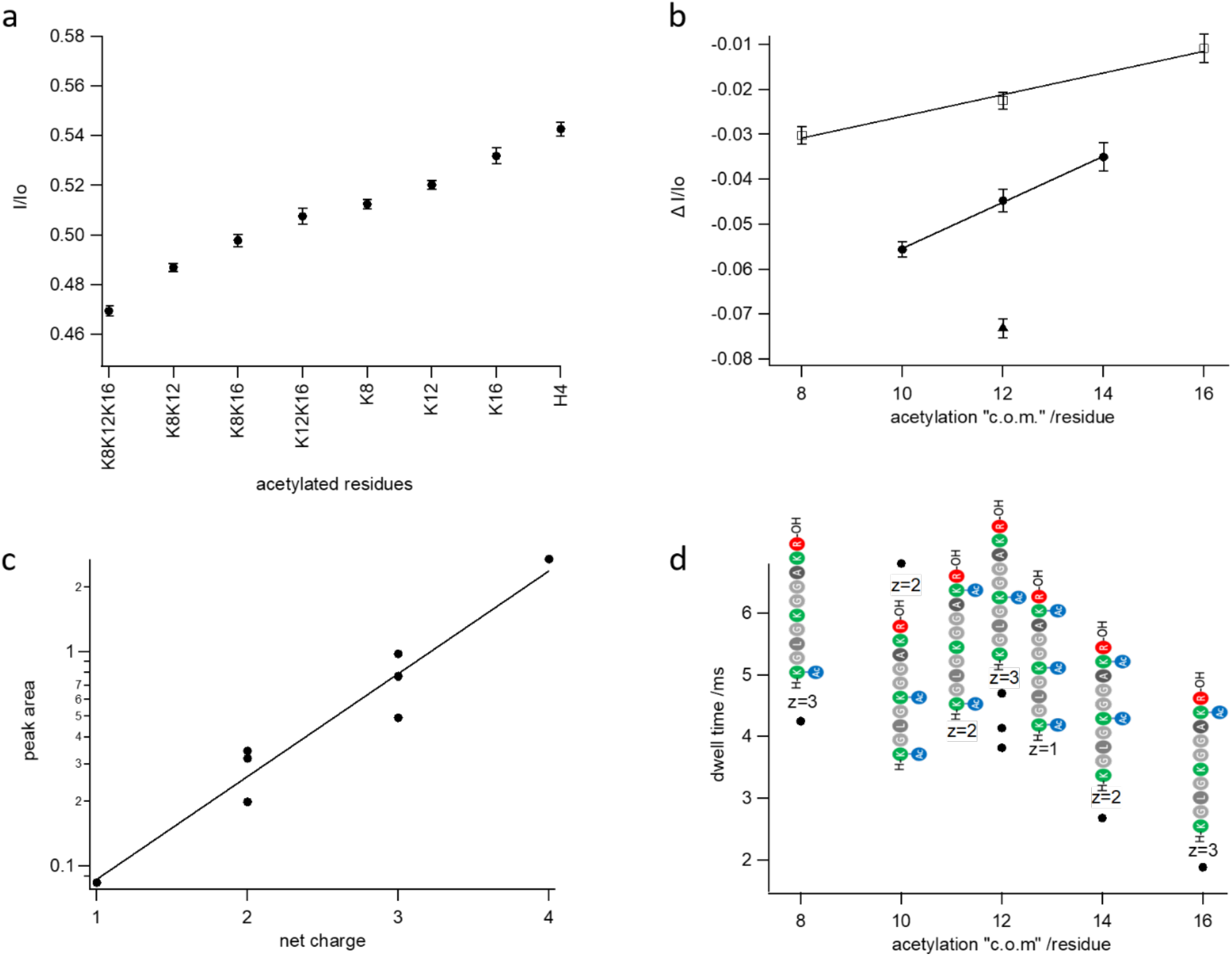
**a**, Positions of maxima for interaction of H4f., H4f.K8Ac, H4f.K12Ac, H4f.K16Ac, H4f.K8K12Ac, H4f.K8K16Ac, H4f.K12K16Ac and H4f.K8K12K16Ac determined with the R220S mutant. Error bars show full width at half maximum of Voigt fits (**Fig. 1**d). **b**, Shift in I/Io produced by acetylation plotted against the mean position or, center of mass, c.o.m.” of the modification for single (open squares), double (filled circles) and triple acetylation (filled triangles). **c**, Logarithmic dependence of peak area as a measure of event frequency on net charge of the peptide. Note decrease of frequency with loss of net charge due to acetylation. **d**, Characteristic dwell-times (see **Extended Data Fig. 4**) vs. acetylation, center of mass”. Note increase from single acetylation at position 16 to double acetylation at positions 8 and 12. Charge numbers are given to show the absence of dependence of dwell-time on peptide charge.

**Extended Data Fig. 4.**
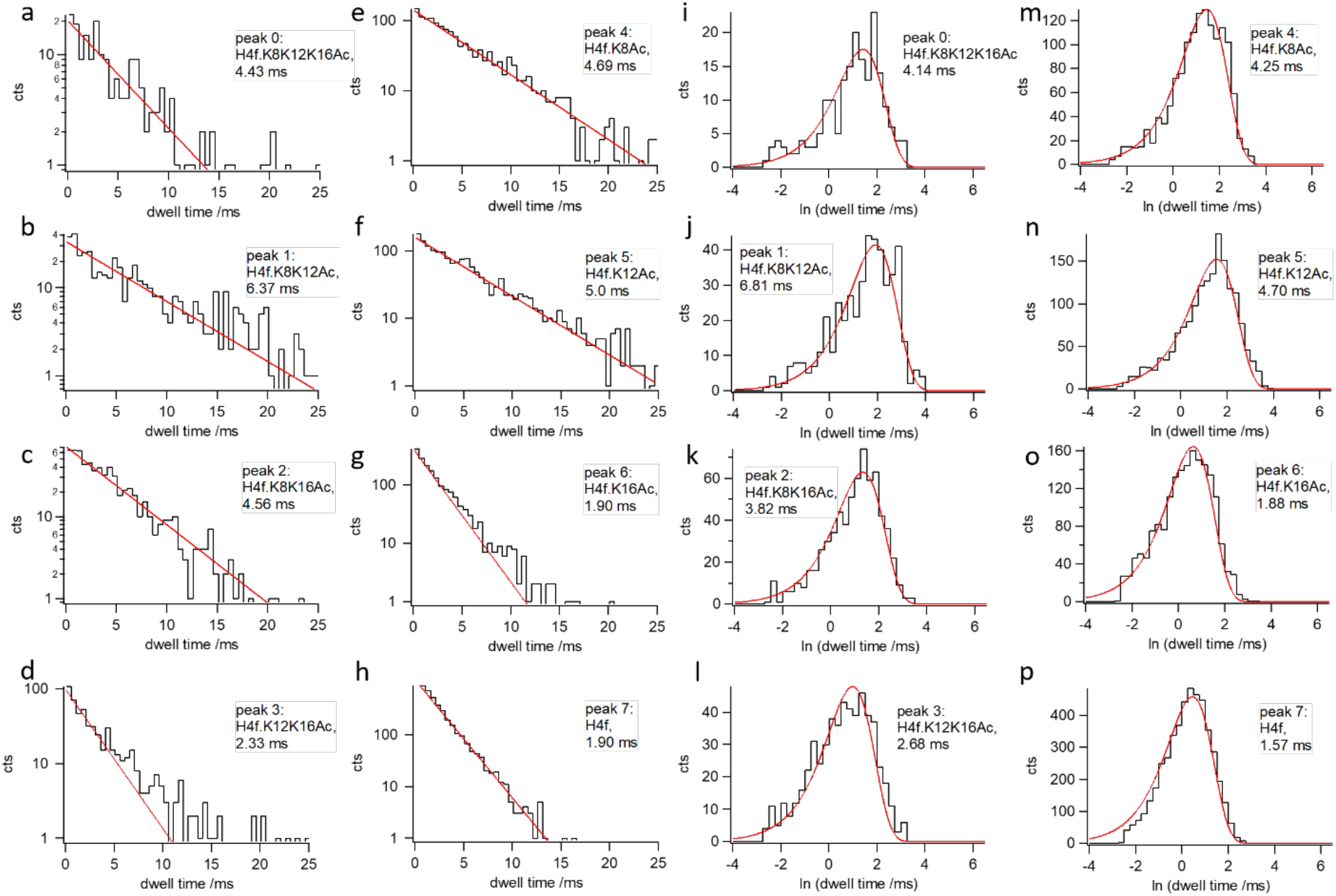
Dwell-time distributions for interaction of H4f., H4f.K8Ac, H4f.K12Ac, H4f.K16Ac, H4f.K8K12Ac, H4f.K8K16Ac, H4f.K12K16Ac and H4f.K8K12K16Ac with the R220S mutant. **a-h**, semilogarithmic plots; **i-p**, histograms of natural logarithm of dwell-times. Red lines: monoexponential fits.

**Extended Data Fig. 5.**
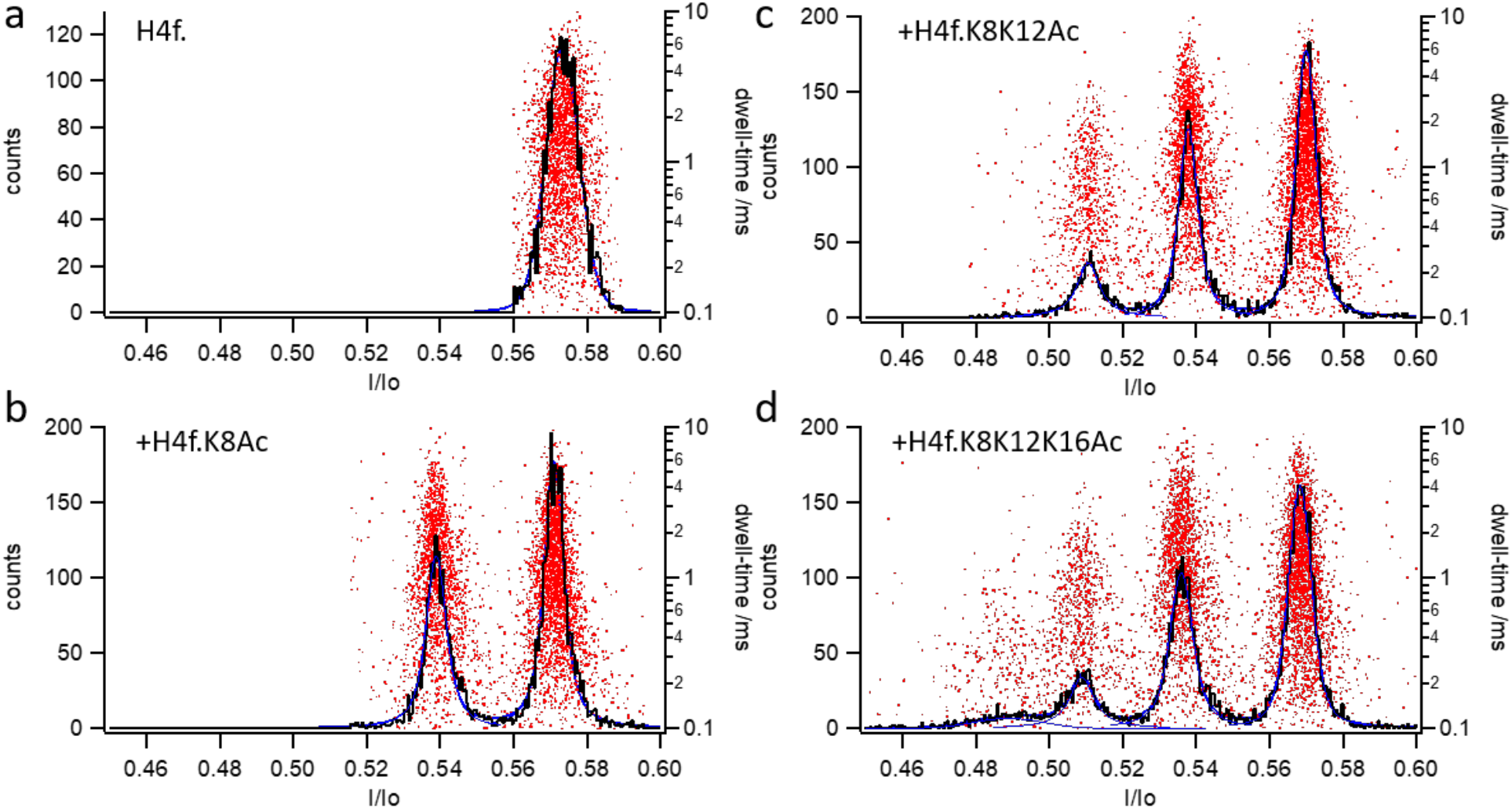
Assignment of maxima to species H4f., H4f.K8Ac, H4f.K8K12Ac and H4f.K8K12K16Ac using sequential peptide addition with the wt-AeL pore.

**Extended Data Fig. 6.**
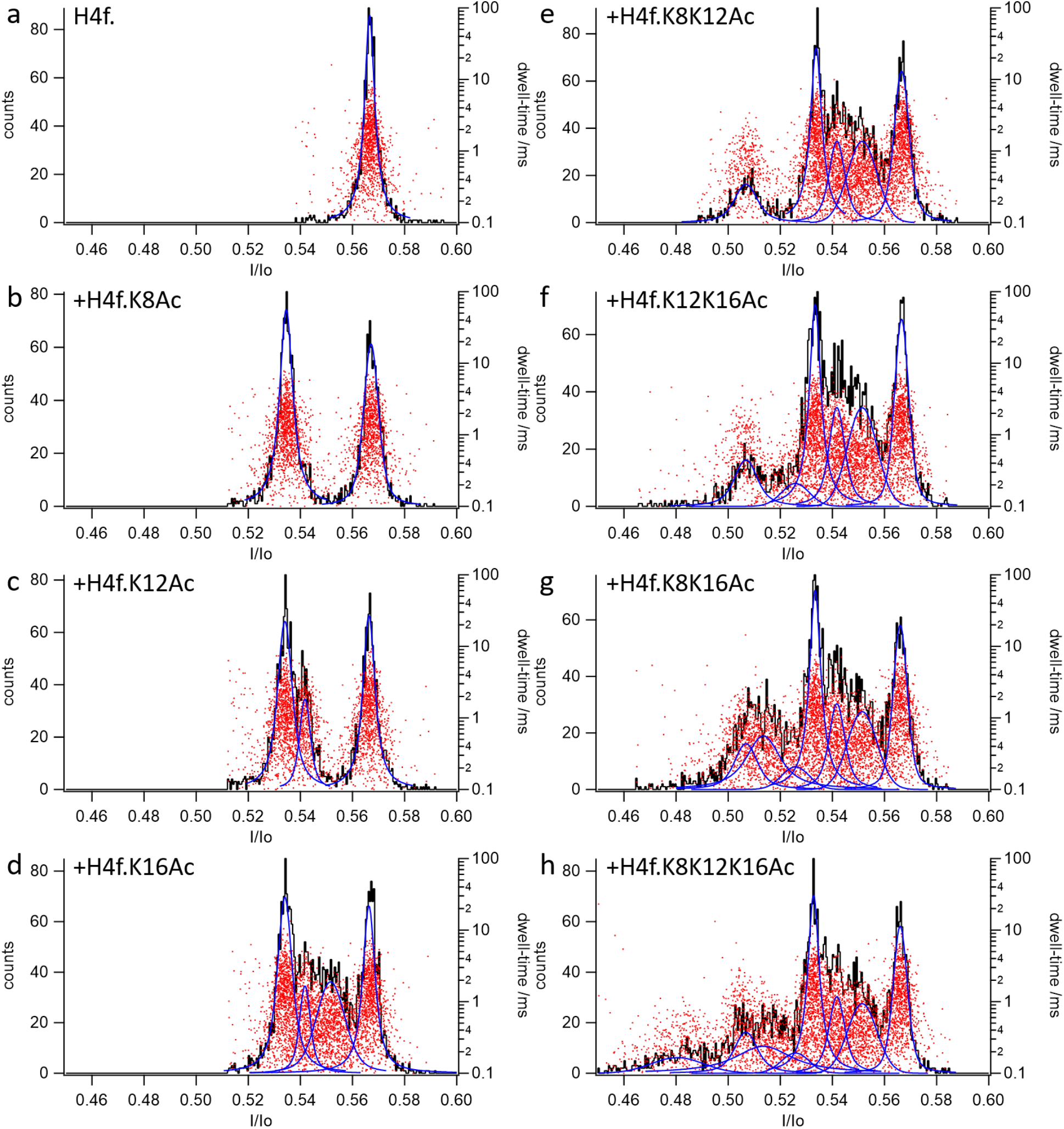
Assignment of maxima to species H4f., H4f.K8Ac, H4f.K12Ac, H4f.K16Ac, H4f.K8K12Ac, H4f.K8K16Ac, H4f.K12K16Ac and H4f.K8K12K16Ac using sequential peptide addition with the wt-AeL pore.

**Extended Data Fig. 7.**
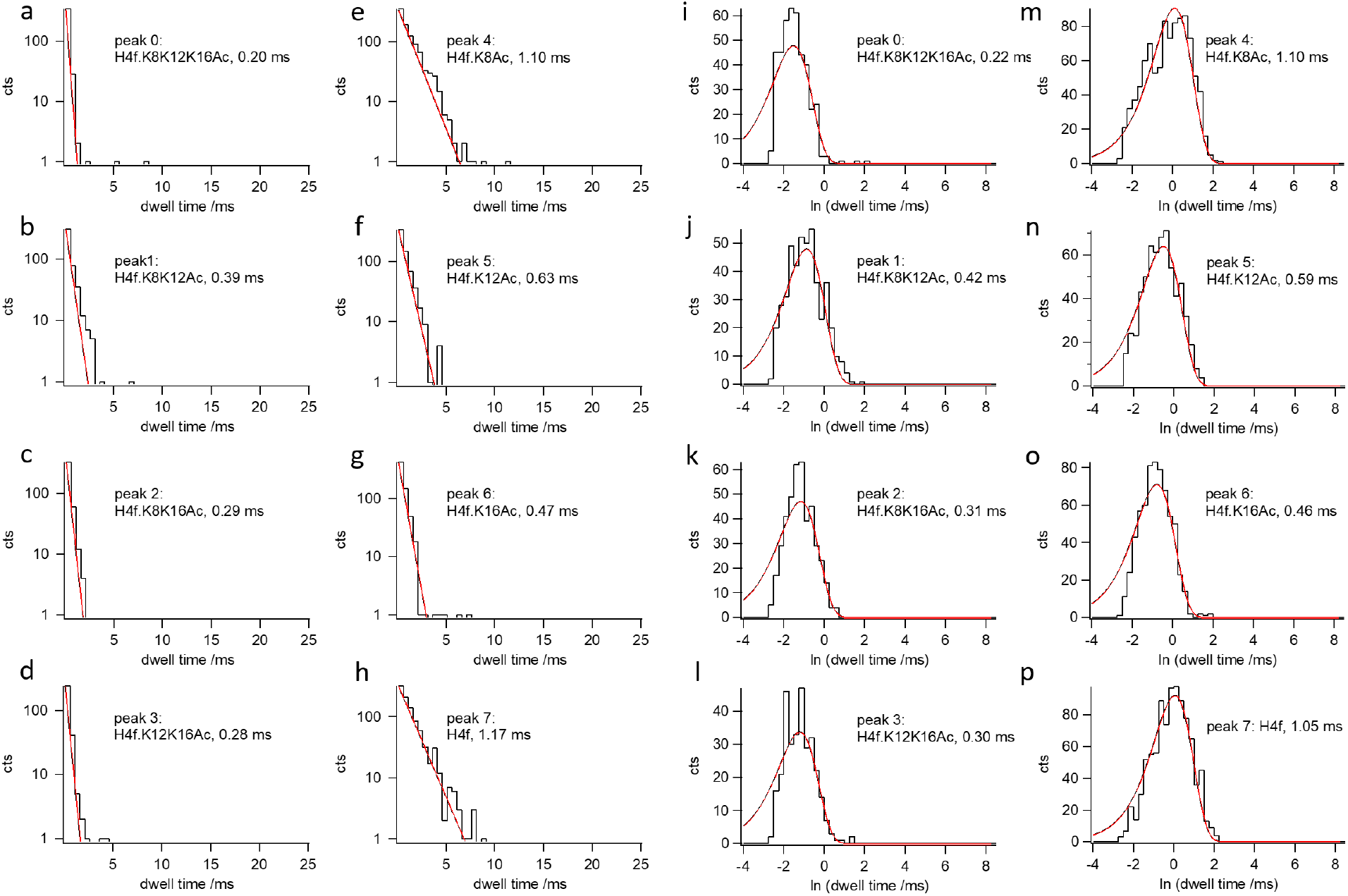
Dwell-time distributions for interaction of H4f., H4f.K8Ac, H4f.K12Ac, H4f.K16Ac, H4f.K8K12Ac, H4f.K8K16Ac, H4f.K12K16Ac and H4f.K8K12K16Ac with the wt-AeL pore. **a-h**, Semilogarithmic plots. **i-p**, Histograms of natural logarithm of dwell-times. Red lines: monoexponential fits.

**Extended Data Fig. 8.**
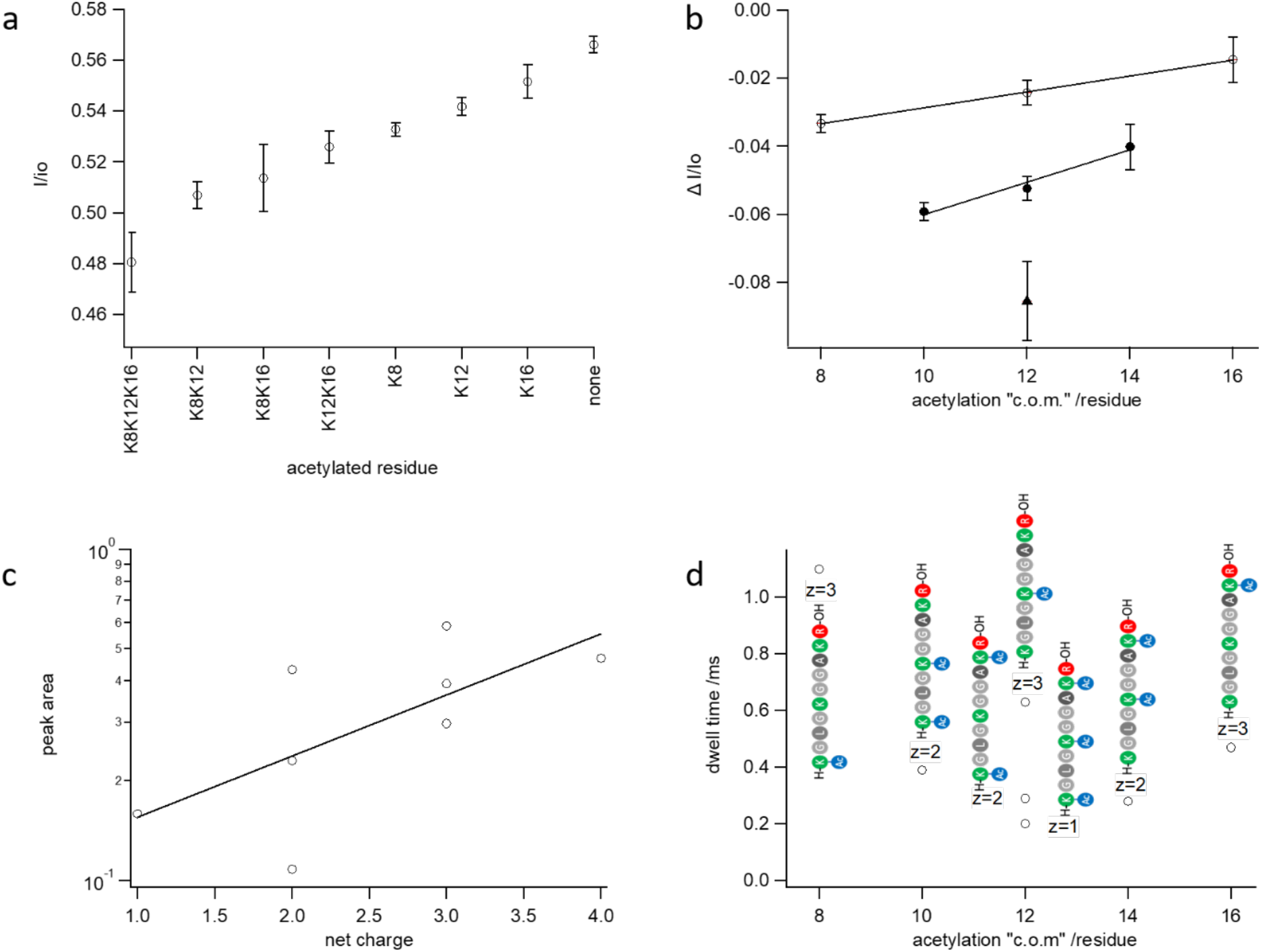
**a**, Positions of maxima for interaction of H4f., H4f.K8Ac, H4f.K12Ac, H4f.K16Ac, H4f.K8K12Ac, H4f.K8K16Ac, H4f.K12K16Ac and H4f.K8K12K16Ac with the wt-AeL pore. Error bars show full width at half maximum of Voigt fits (**Fig. 1**). **b**, Shift in I/Io produced by acetylation plotted against the mean position or, center of mass, c.o.m.” of the modification for single (open squares), double (filled circles), and triple acetylation (filled triangles). **c**, Logarithmic dependence of peak area as a measure of event frequency on net charge of the peptide. Note decrease of frequency with loss of net charge due to acetylation. **d**, Characteristic dwell-times (see **Extended Data Fig. 4**) vs. acetylation, center of mass”. Note decrease from single acetylation at position 8, 12 or 16 to double acetylation at positions 8 and 12 or 8 and 16. Charge numbers are given to show the absence of dependence of dwell-time on peptide charge.

**Extended Data Fig. 9.**
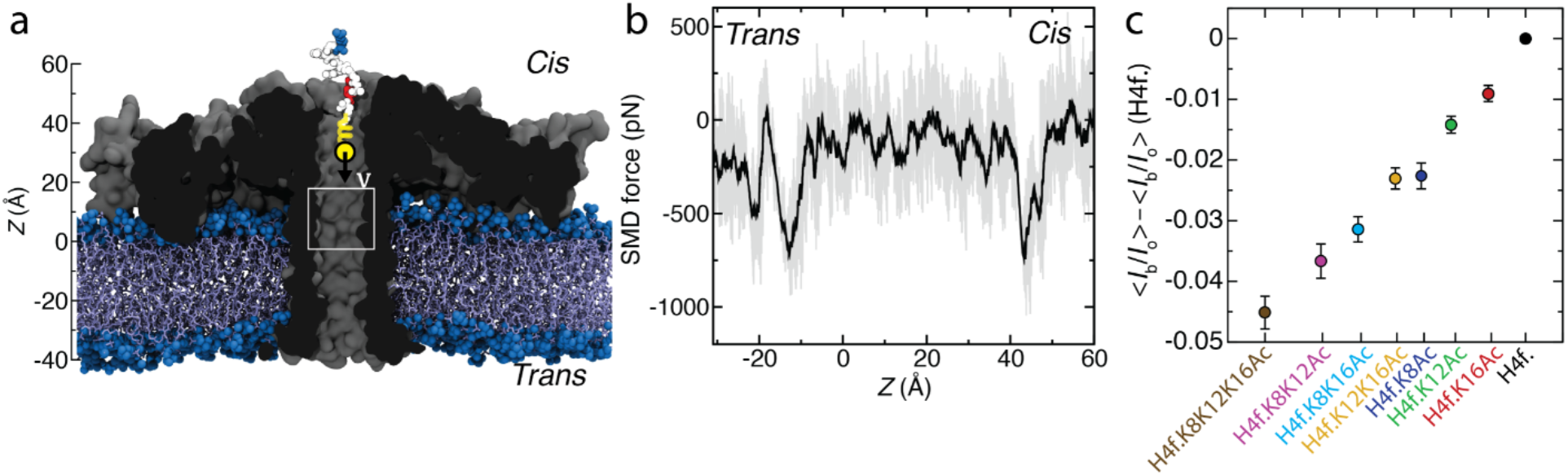
Steered MD simulation of H4f.K8Ac peptide through R220S aerolysin. **a**, initial state of a 100 ns steered molecular dynamics simulation (same as in **Fig. 2**a) where an H4f.K8Ac peptide (vdw spheres) is pulled by a harmonic spring with a constant velocity of 1 Å/ns through an aerolysin nanopore (cutaway molecular surface), embedded in a lipid membrane (blue) and submerged in 2 M KCl electrolyte (not shown). The C-terminus of the peptide is oriented towards the trans-side of the membrane. The white rectangle indicates the region (−2 Å < z < 18 Å) used for averaging in panel c. **b**, Force exerted by the SMD spring (running average: 0.5 Å) vs. the z coordinate of the H4f.K8Ac peptide; **c**, Simulated average relative currents produced by the acetylated peptides calculated as a difference from the baseline H4f. peptide within the region highlighted in the inset of **Fig. 2**b (−2 Å < z < 18 Å). The error bars show the standard error calculated using 80 current values

**Extended Data Fig. 10.**
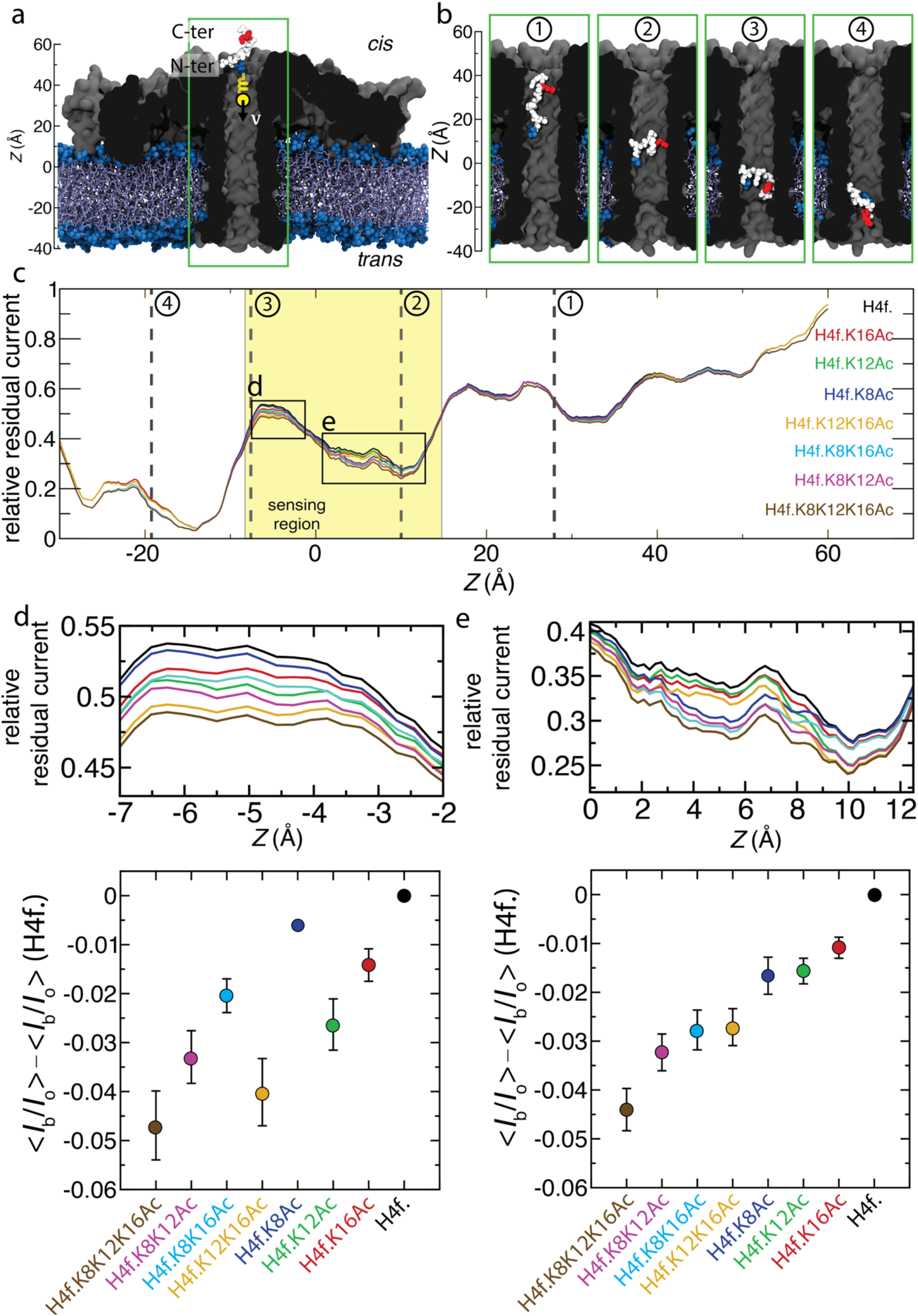
Blockade current of acetylation modification depends on the global conformation of the peptide in the nanopore. **a**, initial state of a 100 ns steered MD simulation where an H4f.K8Ac peptide (vdW spheres) is pulled by a harmonic spring with a constant velocity of 1 Å/ns through an aerolysin nanopore (cutaway molecular surface), embedded in a lipid membrane (blue) and submerged in 2 M KCl electrolyte (not shown). The N-terminus of the peptide is oriented towards the trans-side of the membrane. The orientation is opposite to that shown in **Fig. 2**a. The green box shows the nanopore volume shown in panel b. **b**, Representative conformations of the H4f.K8Ac peptide along the SMD simulation chronologically shown from left to right. As the simulation progresses, the peptide reverses its orientation leading to the C-terminal end of the peptide facing the trans-side of the pore while the peptide exits the pore. **c**, Relative residual current versus center of mass z coordinate of the acetylated H4f. peptides. The coordinate axis is defined in panel a. The conformations from panel b are marked using vertical dotted lines and numbered circles defined in panel b. The currents were computed using SEM ^42^ and were running-averaged with a 5 Å window. The highlighted region shows the sensing volume (−9 Å < z < 15 Å) of the wt-AeL pore ^23^. Two black boxes mark the zoomed in region highlighted in panels d and e. **d**, Relative residual current versus center of mass z coordinate between z=−7 Å and z=−2 Å as highlighted in c. The lower panel shows the average relative residual current and its standard error over the 21 data points in the shown region. The ordering of the peptide currents depends on the peptide conformation. The peptides are arranged in the x-axis in an increasing order according to their experimental blockade values. **e**, Same as panel d but for the region between z=0 Å to z=12.5 Å (53 data points), highlighted in c.

**Extended Data Fig. 11.**
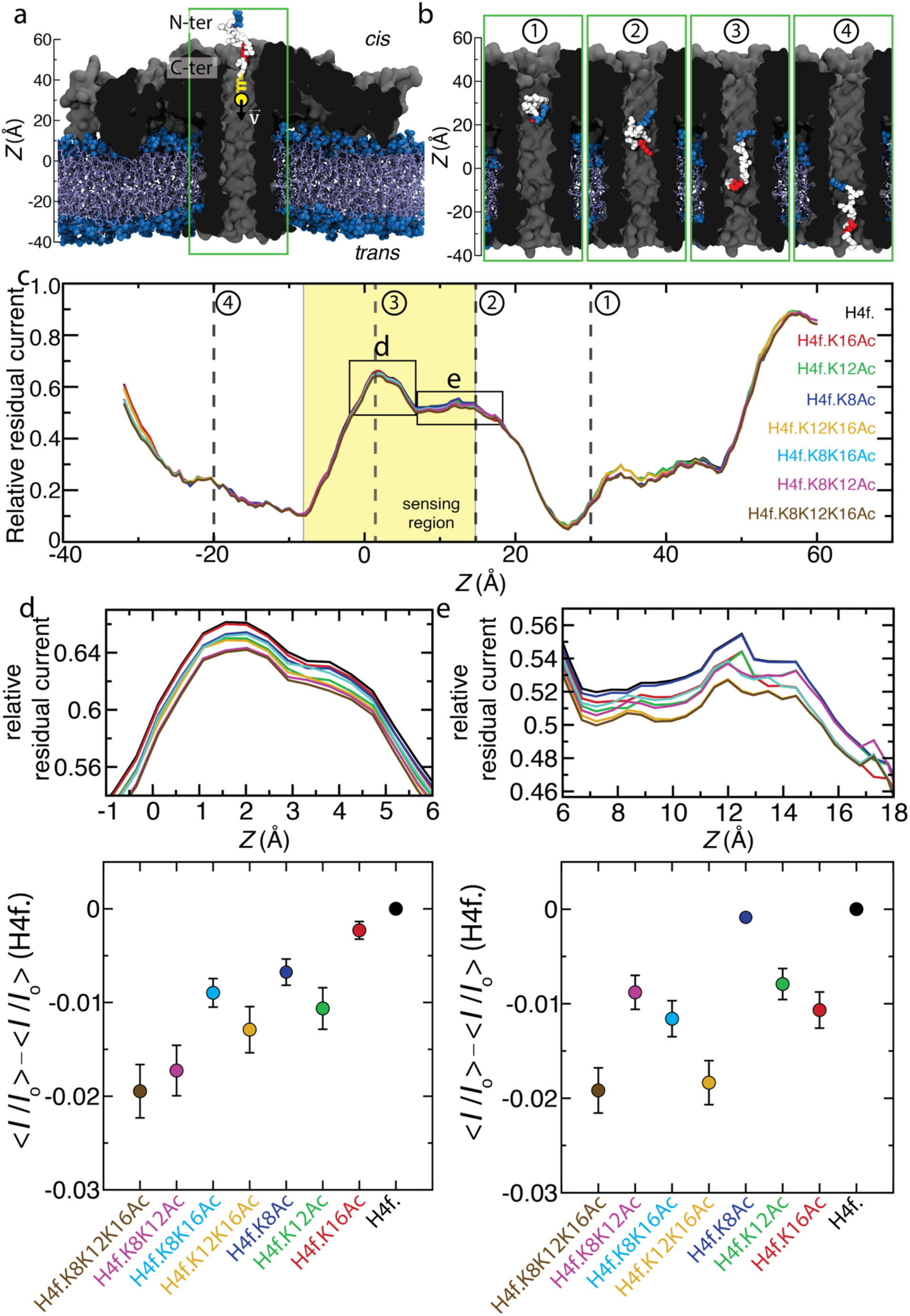
Blockade current of acetylation modification depends on the global conformation of the peptide in the nanopore. **a**, Initial state of a 100 ns steered molecular dynamics simulation where an H4f. K8Ac peptide (vdW spheres) is pulled using a harmonic spring with a constant velocity of 1 Å/ns through an aerolysin nanopore (cutaway molecular surface), embedded in a lipid membrane (blue) and submerged in 2 M KCl electrolyte (not shown). The C-terminus of the peptide is oriented towards the trans-side of the membrane. The global orientation of the peptide is the same as that in the trajectory shown in **Fig. 2**a, but the simulation stars from a slightly different microscopic configuration. The green box shows the nanopore volume shown in panel b. **b**, Representative conformations of the H4f.K8Ac peptide along the SMD simulation chronologically shown from left to right. **c**, Relative residual current versus center of mass z coordinate of the acetylated H4f. peptides. The coordinate axis is defined in panel a. The conformations from panel b are marked using vertical dotted lines and numbered circles defined in panel b. The currents were computed using SEM ^42^ and were running-averaged with a 5 Å window. The highlighted region shows the sensing volume (-9 Å < z < 15 Å) of the wt-AeL pore ^23^. Two black boxes mark the zoomed in region highlighted in panels d and e. **d**, Relative residual current versus center of mass z coordinate between z=-1 Å and z=6 Å as highlighted in c. The lower panel shows the average relative residual current and its standard error over the 15 data points in the shown region. The ordering of the peptide currents depends on the peptide conformation. The peptides are arranged in the x-axis in an increasing order according to their experimental blockade values. **e**, Same as panel d but for the region between z=6 Å to z=18 Å (24 data points), highlighted in c.

**Extended Data Fig. 12.**
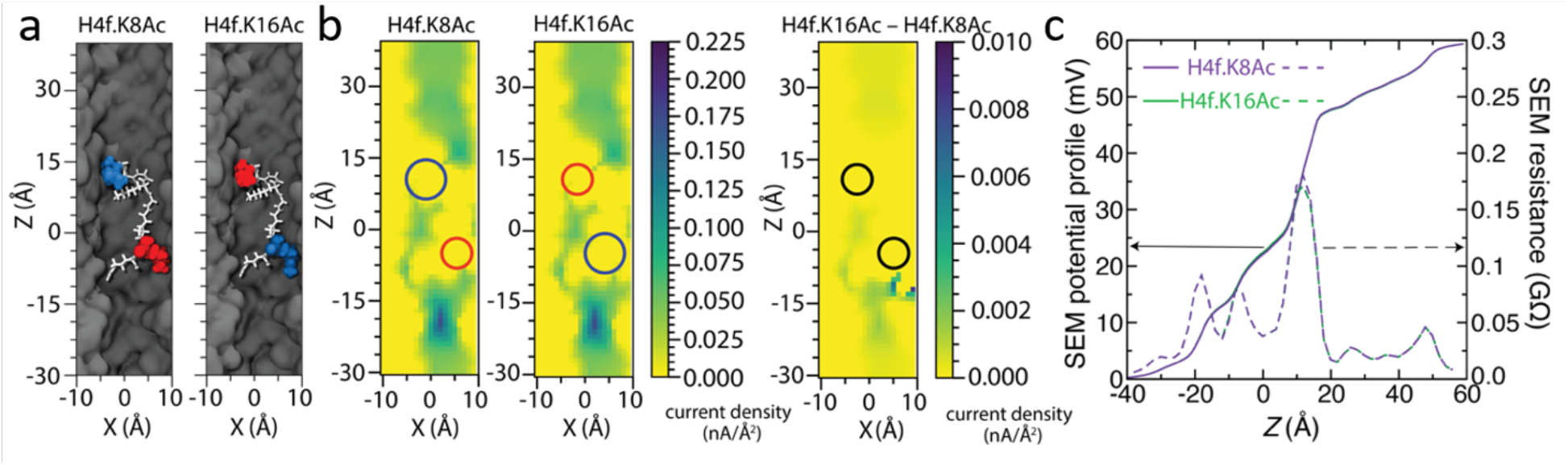
SEM analysis of H4f.K8Ac vs. H4f.K16Ac for another conformation of the H4f. peptide taken from the same SMD trajectory as that shown in Fig 2 of the main text. **a**, Representative conformation (z height of 5 Å) of a H4f. K8Ac peptide in the sensing region of the nanopore and the corresponding computational model of the H4f.K16Ac peptide. The peptide residues are drawn as white sticks except the K8 and K16 residues which are shown as vdW spheres colored according to their acetylation state: acetylated (blue) and unmodified (red). **b**, Local density of transmembrane ion current (its z component) near the H4f.K8Ac (left) and H4f.K16Ac (center) peptides and their difference (right). The currents were computed using SEM over a 1 Å grid. The heat map shows a cross section of the nanopore volume along the pore axis. The circles indicate the approximate locations of the K8 and K16 residues. **c**, Electrostatic potential along the nanopore axis for the H4f.K8Ac (purple) and H4f.K16Ac (green) peptide systems (solid lines, left axis) and local resistance of the nanopore volume (dashed lines, right axis). The local resistance was computed from the local electrostatic potential using 4 Å segments.

**Extended Data Fig. 13.**
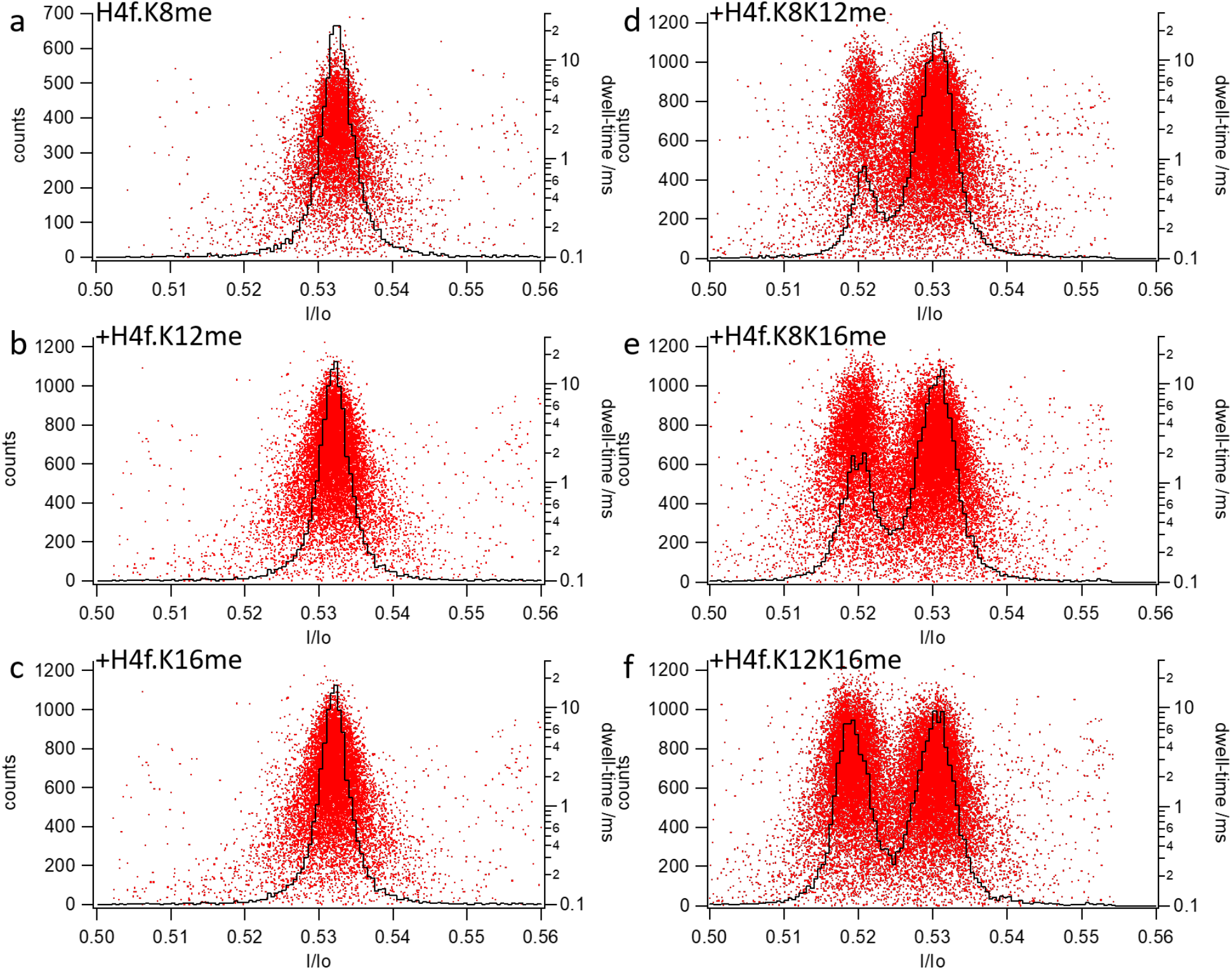
Assignment of maxima for singly and doubly monomethylated peptoforms of H4f. using sequnetianl addition with the R220S mutant.

**Extended Data Fig. 14.**
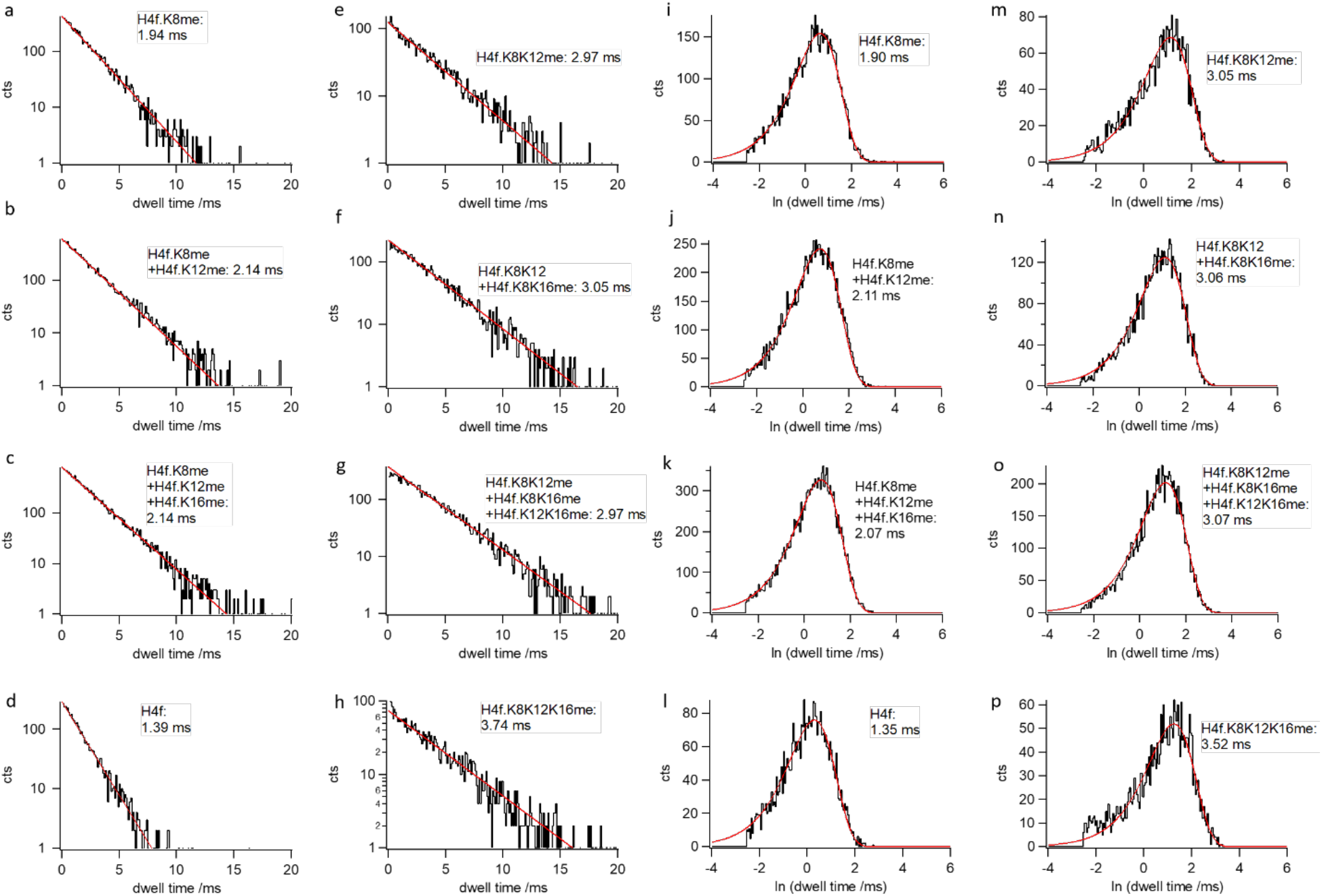
Dwell-time distributions for the interaction of singly, doubly and triply monomethylated peptoforms of H4f with the R220S mutant.

**Extended Data Fig. 15.**
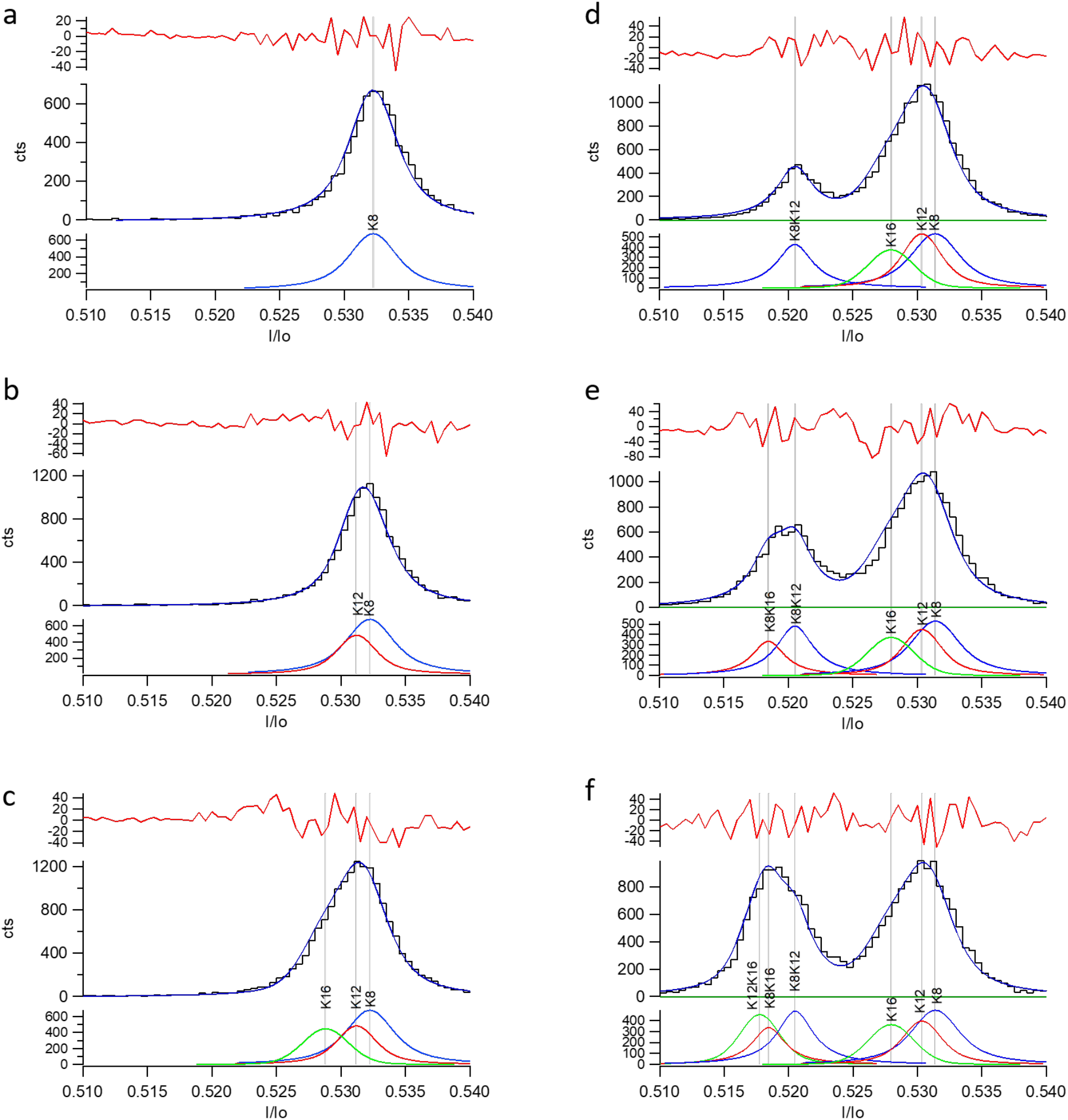
discrimination of singly (**a-c**) and doubly (**d-f**) monomethylated isomeric H4f. peptoforms using sequential addition with the R2020S mutant.

**Extended Data Fig. 16.**
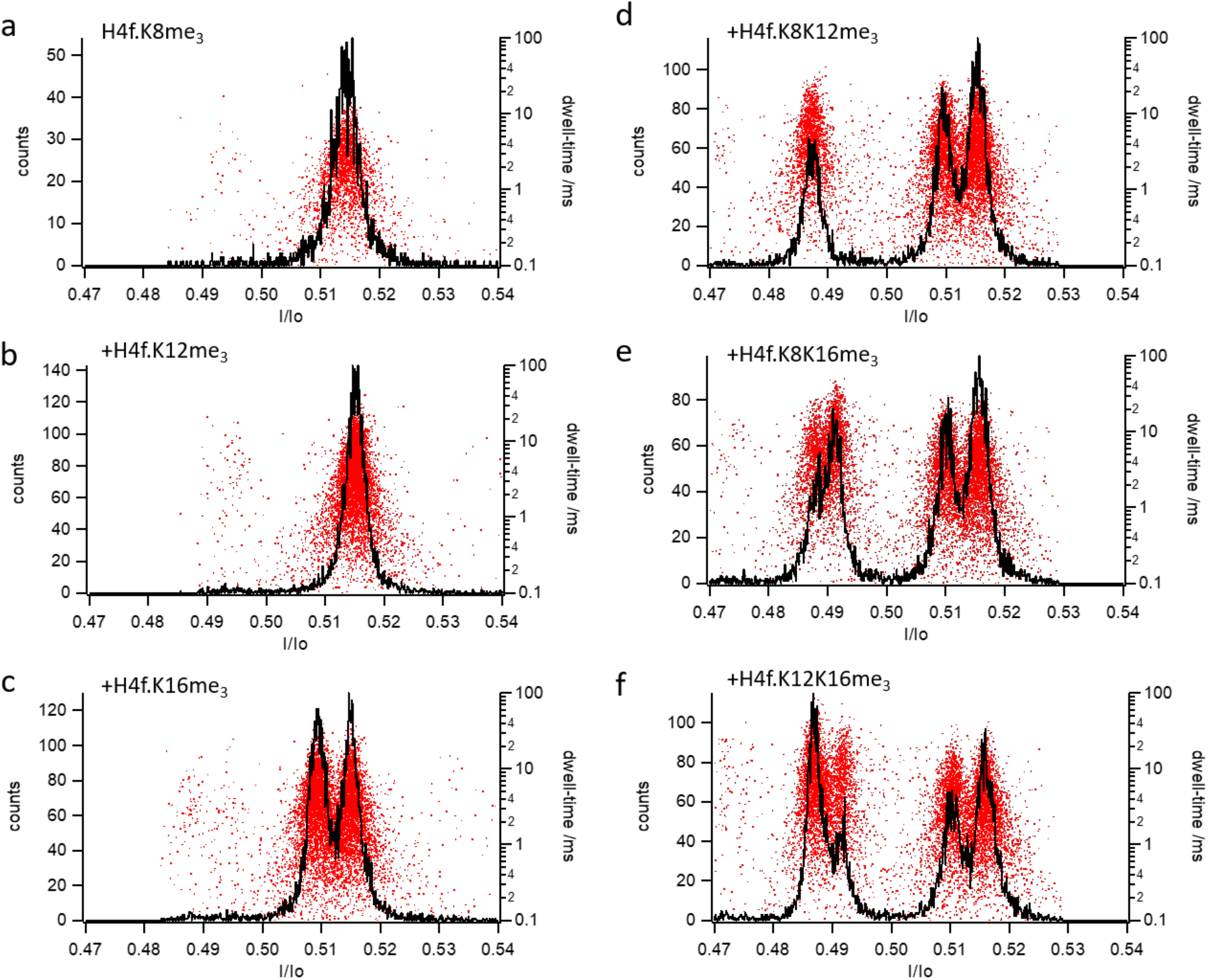
Assignment of maxima for singly and doubly trimethylated peptoforms of H4f. using sequential addition with the R220S mutant.

**Extended Data Fig. 17.**
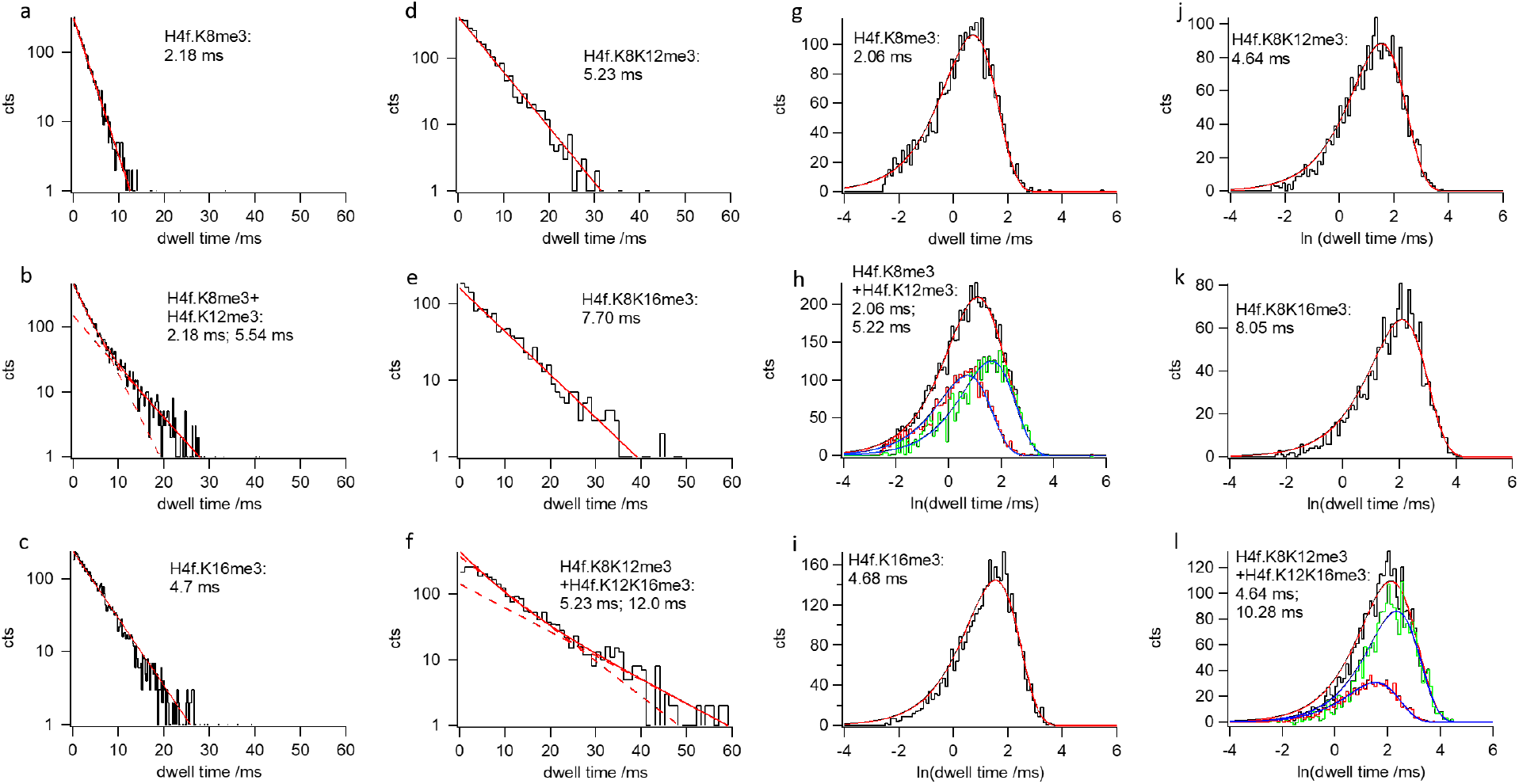
Dwell-time distributions for the interaction of singly, doubly and triply trimethylated peptoforms of H4f. with the R220S mutant.

**Extended Data Fig. 18.**
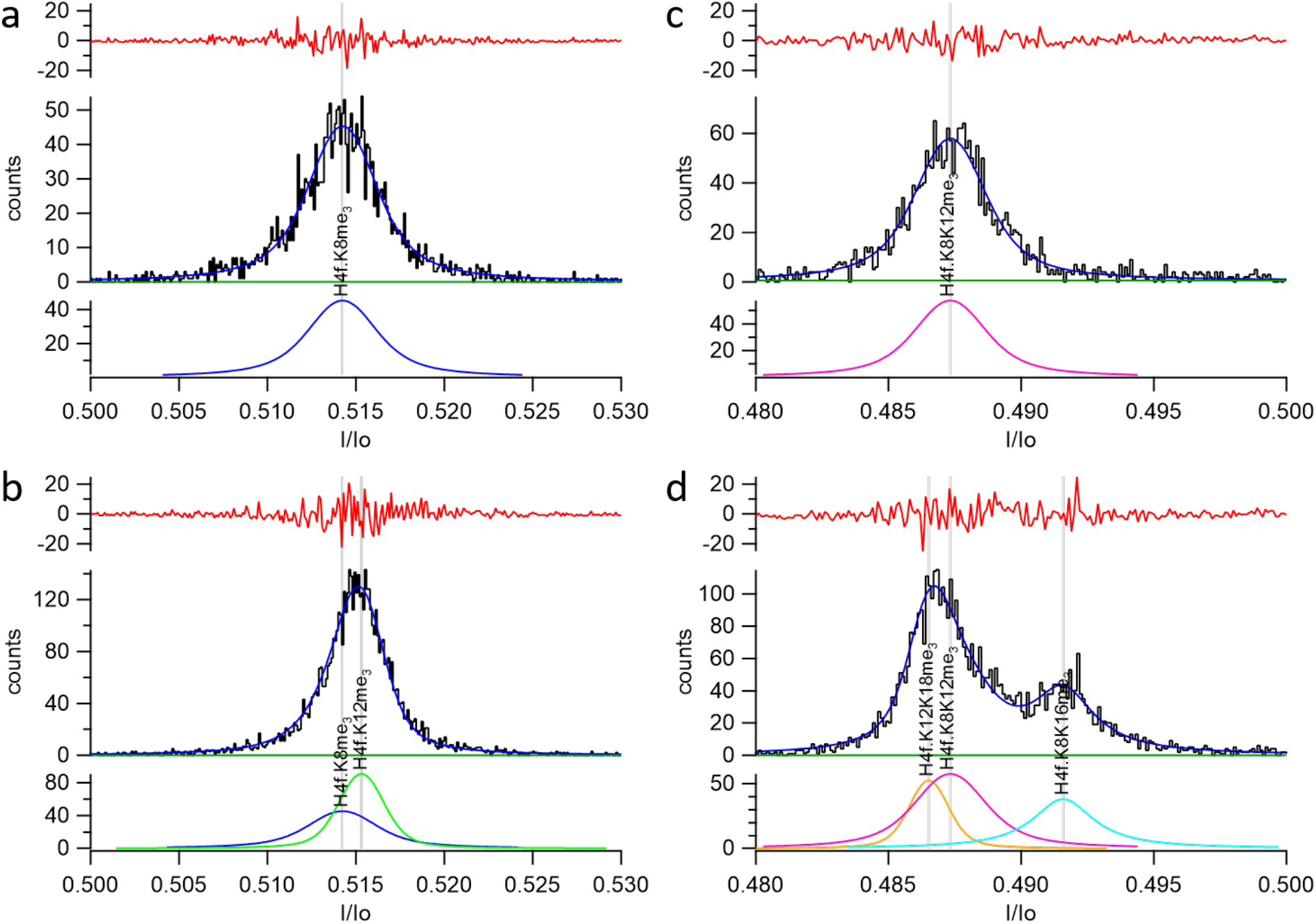
Discrimination of singly (**a-c**) and doubly (**d-f**) trimethylated isomeric H4f. peptoforms with the R220S mutant.

**Extended Data Fig. 19.**
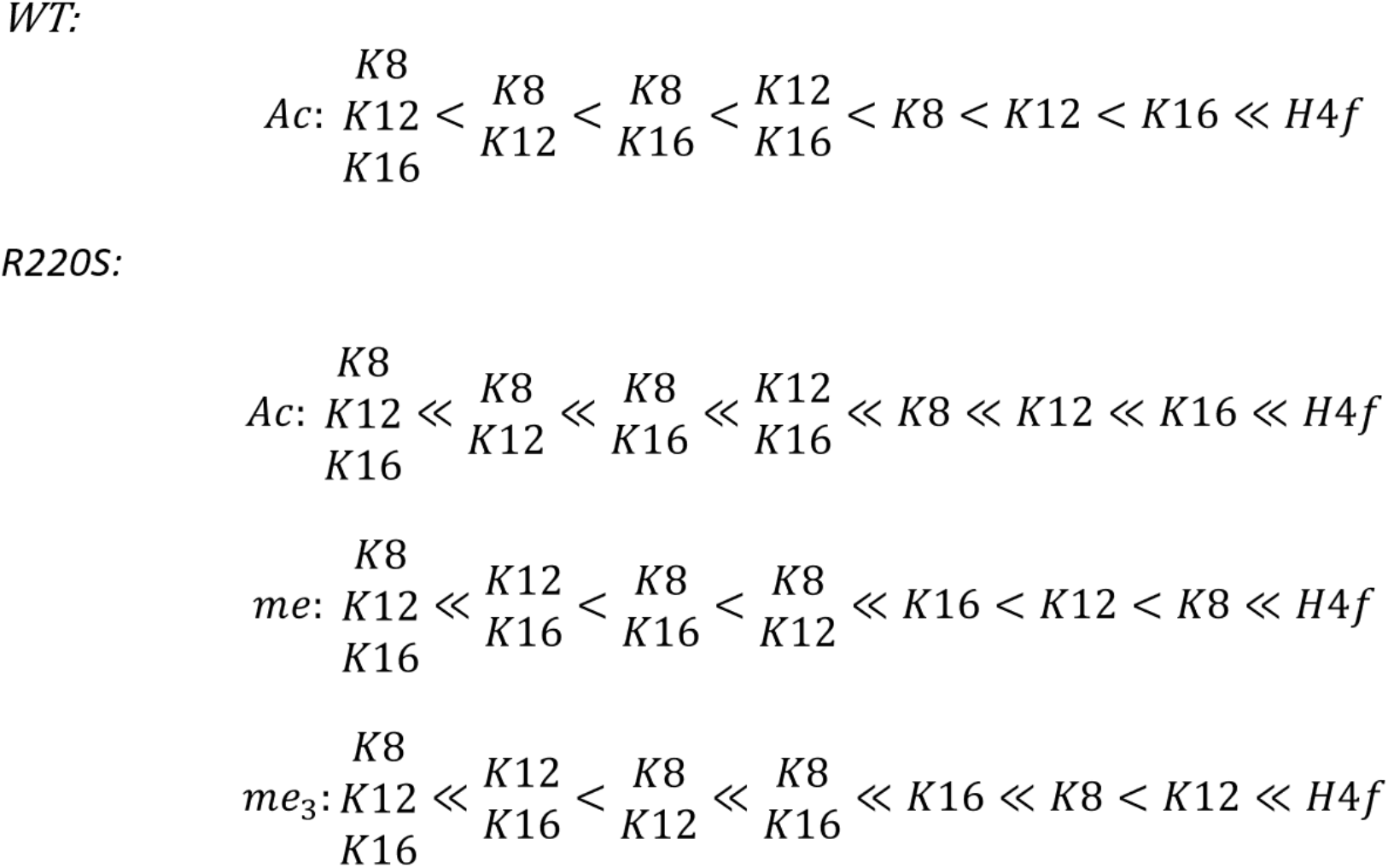
Overview of the sequences of depth of block (I/Io) observed in this study.

**Extended Data Fig. 20.**
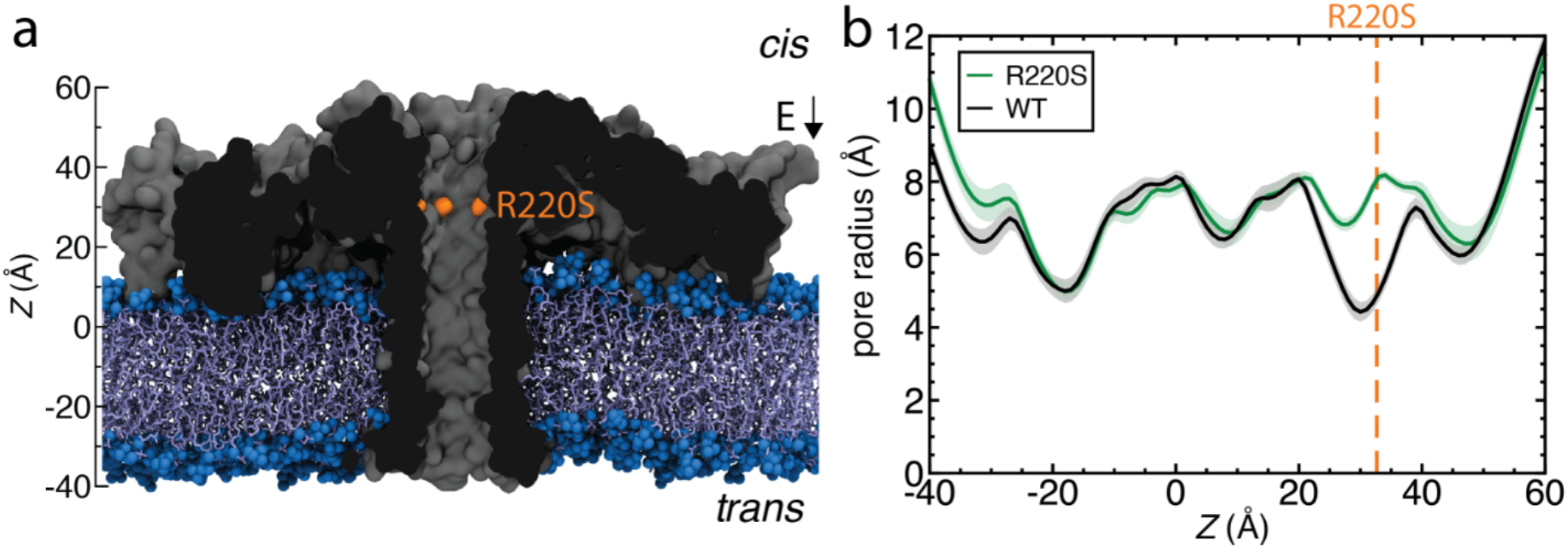
Structure of R220S mutant aerolysin. **a**, Initial state of an MD simulation where a R220S aerolysin nanopore (cutaway molecular surface) is embedded in a lipid membrane (blue), submerged in 2 M KCl electrolyte (not shown) and simulated under an applied electric field corresponding to a transmembrane voltage bias of −60 mV, as used experimentally in this study Residue R220 is shown in orange. **b**, Pore radius of the R220S pore (green) and the wt-AeL pore (black)^23^ is plotted along the symmetry axis of the pore. The radius is calculated by using the HOLE software and averaged over the last 20 ns of the MD simulation trajectory for R220S and wt-AeL nanopore.

**Extended Data Fig. 21.**
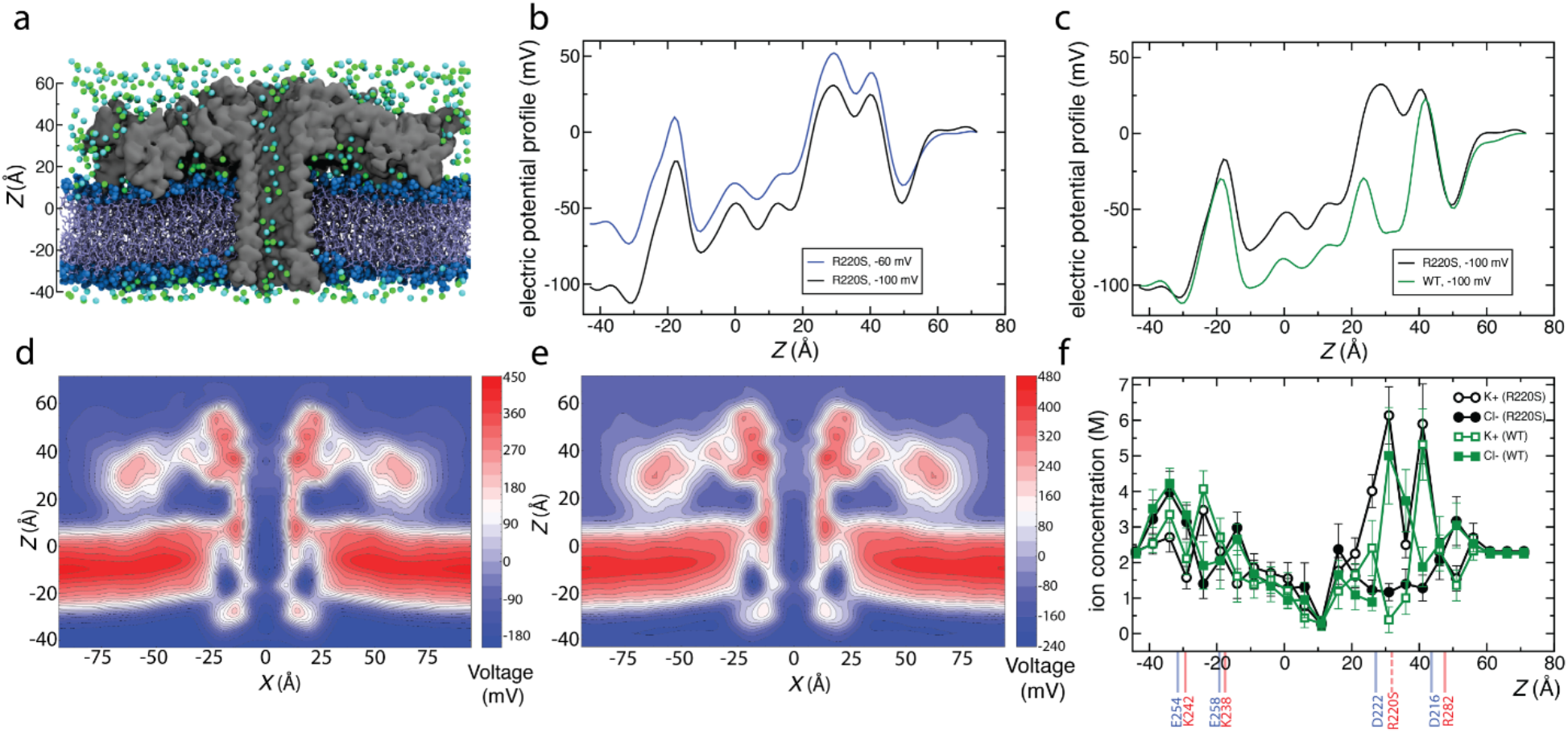
Ion transport driven by electric field through R220S and wt-AeL nanopore. **a**, Initial state of an MD simulation where a R220S aerolysin nanopore (gray cutaway molecular surface) is embedded in a lipid membrane (blue), submerged in 2 M KCl electrolyte (potassium and chloride shown as cyan and green spheres respectively) and simulated under an applied electric field corresponding to a transmembrane voltage bias of −60 mV, as used experimentally in this study. **b**, average electrostatic potential along the symmetry axis of R220S aerolysin (the z axis) at −60 mV (blue) and −100 mV (black). The plot was obtained by taking the average of instantaneous values of electrostatic potential along the symmetry axis over a 70 ns (R220S) and a 90 ns (wt) MD trajectory; **c:**average electrostatic potential along the symmetry axis of R220S aerolysin (the z axis) at −100 mV (black) and wt-AeL at −100 mV and 2 M KCl electrolyte (green). The plot of R220S and wt-AeL was obtained by taking the average of instantaneous values of electrostatic potential along the symmetry axis over a 70 ns and 90 ns MD trajectory respectively. The average MD current is −0.13 nA at −100 mV for the wt-AeL pore and −0.21 nA at −100 mV for the R220S aerolysin pore. **d, e**, Two-dimensional electrostatic potential map of R220S aerolysin at −60 mV (d) and −100 mV (e) transmembrane bias. The map was obtained by averaging instantaneous distributions of electrostatic potentials over a 70 ns (R220S) and 90 ns (wt) MD trajectory and the sixfold symmetry of the channel. **f**, profiles of potassium (black, open circles) and chloride (black, filled circles) ion concentration along the symmetry axis of the R220S aerolysin nanopore. Also shown are the profiles of potassium (green, open squares) and chloride (green, filled squares) ion concentration along the symmetry axis of the wt-AeL nanopore. The ion concentration values are averaged over 5 Å z bins inside the pore (defined by the radius profile shown in **Extended Data Fig. 20**b) every 0.25 ns and further block averaged over 10 ns intervals in a total simulation time of 70 and 90 ns for the R220S and wt-AeL nanopore respectively. The ion concentration for z values outside the protein are calculated by counting the potassium and chloride ions in water molecules outside the protein that constitute the bulk water in the simulation system. Background blue and red lines represent positive and negative amino acid residues along the inner surface of the aerolysin pore, which are annotated by the amino acid type and residue number at the bottom. The dashed blue line represents the R220 positive amino acid that is mutated to a neutral serine in the R220S aerolysin pore.

